# MPSeqM, a tool combining multiplex PCR and high-throughput sequencing to study the polymorphism of eight *Leptosphaeria maculans* avirulence genes and its application to field surveys in France

**DOI:** 10.1101/2023.10.06.561155

**Authors:** Angélique Gautier, Valérie Laval, Marie-Hélène Balesdent

## Abstract

*Leptosphaeria maculans* is a fungal pathogen causing stem canker of oilseed rape (*Brassica napus*). The disease is mainly controlled by the deployment of varieties with major resistance genes (*Rlm*). *Rlm* genes can rapidly become ineffective following the selection of virulent isolates of the fungus, i.e. with deletions or mutations in the corresponding avirulence genes (*AvrLm*). Reasoned and durable management of *Rlm* genes relies on the detection and monitoring of virulent isolates in field populations. Based on previous knowledge of *AvrLm* gene polymorphism, we developed a tool combining multiplex PCR and Illumina sequencing to characterise allelic variants for eight *AvrLm* genes in field *L. maculans* populations.

We tested the method on DNA pools of 71 characterised *L. maculans* isolates and of leaf spots from 32 *L. maculans* isolates. After multiplex-PCR and sequencing with MiSeq technology, reads were mapped on an in-house *AvrLm* sequence database. Data were filtered using thresholds defined from control samples included in each run. Proportions of each allelic variant per gene, including deletions, perfectly correlated with expected ones. The method was then applied to around 1300 symptoms (42 pools of mainly 32 leaf spots) from nine *B. napus* fields. The proportions of virulent isolates estimated by sequencing leaf spot pools perfectly correlated with those estimated by pathotyping. In addition, the proportions of allelic variants determined at the national scale also correlated with those previously determined following individual sequencing of *AvrLm* genes in a representative collection of isolates. Finally, the method also allowed us to detect still undescribed and rare allelic variants. Despite the diversity of mechanisms generating virulent isolates and the gene-dependant diversity of *AvrLm* gene polymorphism, the method proved suitable for large-scale and regular monitoring of *L. maculans* populations, which will make it possible to choose effective *Rlm* genes and to detect resistance breakdowns at early stages.

## INTRODUCTION

The fight against crop pests is a major issue for food safety in a changing environmental and socio-economic context. Whether chemical or genetic, control measures are more effective as they are adapted to local pest populations. These populations also are constantly evolving in response to the deployment of chemical or genetic protection solutions. For instance, pesticide-resistant individuals regularly appear in fungal or insect populations following the large-scale use of pesticides (McCulloch & Waters, 2023). The selection of such individuals in populations renders the pesticides ineffective. Similarly, plant resistance genes introduced in crop varieties and effective against fungal isolates possessing matching avirulence genes exert strong selection pressure on pathogenic populations. The selection of virulent isolates in populations rapidly decreases the efficacy of the genetic solution. Monitoring pest populations and their characterisation regarding their potential resistance to control measures is thus a main issue, not only in choosing an effective management strategy but also in detecting outbreak events and preventing their spread.

Monitoring virulent isolates is particularly important for the fungal ascomycete *Leptosphaeria maculans*, responsible for the stem canker disease of oilseed rape (*Brassica napus*). The main control measure deployed to limit the impact of the disease is genetic control, including the use of specific resistance (*Rlm*) genes. *L. maculans* populations have the potential to evolve rapidly toward virulence, which has been reported in many instances (Rouxel et al., 2003; Sprague et al., 2006, Zhang et al., 2016). Lastly, potentially effective resistance sources are scarce in oilseed rape and the few currently available *Rlm* genes must be carefully managed to avoid new breakdowns (Borhan et al., 2022; Rouxel and Balesdent, 2017). Until recently, the monitoring of *L. maculans* populations consisted of large-scale isolate samplings from diseased *B. napus* plants, followed by inoculation of each isolate on a plant differential set with defined *Rlm* genes (for instance, Alnajar et al., 2022; Balesdent et al., 2006, 2023; Stachowiak et al., 2006; Zhang et al., 2016). These biological tests are however inadequate for high-throughput analyses and for the detection of rare mutation events in populations. They require adapted facilities and materials (seeds of the plant differential set, control isolates, growth chambers with tight temperature and humidity regulation), and they are time- and labor-intensive and thus, costly. Thus, only a limited number of isolates from a limited number of sites can be analysed each year, and the survey results are obtained six to eight months later. In France, large-scale surveys were done on eight to 12 sites at about 10-year intervals (Balesdent et al., 2023). One full working year is needed to fully characterize populations of around 80 isolates per site. The size of such samplings limits the precision of the estimates of virulent isolates in a population. For instance, in a sampling of 80 isolates per site, the detection of five virulent isolates (6.2%) only allows us to say that the actual proportion of virulent isolates ranges between 2% and 14% (<u>Exact CI (inrae.fr)</u>), corresponding to very distinct breakdown situations.

Fortunately, *L. maculans* is a fungus for which we have a detailed knowledge of the genetic determinants involved in the recognition of the fungus by the plant, i.e., its avirulence (*AvrLm*) genes. To date, 11 *AvrLm* genes have been cloned (Borhan et al., 2022), and the molecular mechanisms by which these genes are mutated or inactivated to avoid being recognized by the corresponding *Rlm* genes were analysed in detail for a few avirulence genes, such as *AvrLm3* and *AvrLm4-7* (Balesdent et al., 2022). The study of the *AvrLm* gene polymorphism in a French population revealed contrasting situations, depending on the gene, as regards the level of polymorphism in virulent or avirulent isolates and the type of molecular events responsible for the gain of virulence (Gautier et al., 2023). Some *AvrLm* genes were nearly monomorphic while others were highly polymorphic, and the mutational events in virulent isolates included single nucleotide mutation (SNP), inactivating mutations introducing stop codons, particularly following Reaped-Induced Point mutations (RIP; Fudal et al. 2009; Gladyshev, 2017), deletion of the whole gene including the surrounding region, or the masking of the avirulence function by the presence of another avirulence gene. For instance, the presence of at least some alleles of *AvrLm4-7* is sufficient to mask the *AvrLm3-Rlm3* or the *AvrLm9*-*Rlm9* interactions (Plissonneau et al., 2017, Ghanbarnia et al., 2018). With this knowledge, one can now envisage developing molecular tools for the survey of virulent isolates at one or more avirulence loci.

Molecular methods were already developed for this purpose, but they are often based on unsatisfactory compromises. For instance, some tools only rely on PCR amplification of the avirulence gene, and the percentage of virulent isolates is deduced from the percentage of isolates that fail to amplify the gene. This approach works well for *AvrLm* genes having gene deletion as the main event in virulent isolates, such as *AvrLm1* (Gout et al., 2007). Even though a few virulent isolates display inactivated versions of *AvrLm1*, still amplified in these isolates, these events are rare, with only one such isolate among 89 in the French population (Gautier et al., 2023). The lack of PCR amplification could thus be a good proxy for the frequency of virulent *avrLm1* isolates. A quantitative PCR assay was developed to quantify the proportion of deleted isolates for *AvrLm1*, relative to the ITS (Van de Wouw et al., 2010). In most cases however, the deletion is not the only mutational event in virulent isolates, and deletions may be rare or even never observed for some *AvrLm* genes (Gautier et al., 2023, Van de Wouw et al., 2023).

Thus, *AvrLm* sequencing in individual isolates is the easiest way to access the genotype and to infer its phenotype, provided the correspondence between genotype and phenotype is available. In the situation of epistatic interactions between avirulence genes, such as the masking effect of *AvrLm4-7* on the *AvrLm3*/*Rlm3* and *AvrLm9*/*Rlm9* interactions, the sequencing of the two genes involved in the epistatic interaction should also give access to the phenotype thanks to the detailed work already published on these complex interactions.

Different approaches have been proposed to avoid PCR amplification and sequencing of *AvrLm* genes. Van de Wouw and Howlett (2014) developed a pyrosequencing method to quantify the proportions of virulent isolates toward *Rlm4*. This quantitative method works on pools of individuals and tags one polymorphic site in *AvrLm4-7* known to be responsible for avirulence towards *Rlm4* (Parlange et al., 2009). However, this method gives sequence information on a small part of the gene only (90 bp) and thus cannot be used to infer virulence on *Rlm7*, due to the diversity of mutations responsible for this phenotype (Daverdin et al, 2012; Balesdent et al., 2022). An HRM (High-Resolution Melting analysis) method was also developed to detect sequence polymorphism in *AvrLm4-7* and determine virulence toward *Rlm4* and *Rlm7*. The method worked well in the situation of limited polymorphism of the gene (Carpezat et al., 2014). However, with the recent large-scale use of *Rlm7,* at least in France, and the diversification of sequence polymorphism in virulent isolates (Daverdin et al., 2012; Balesdent et al., 2022), it became more and more complex to convert new HRM profiles into virulent or avirulent alleles.

Finally, whole genome sequencing (WGS) of individual isolates gives direct access to the sequences of the full set of *AvrLm* genes in an isolate. For instance, Chen et al. (2020) studied the polymorphism in nine avirulence genes following the WGS of 162 Canadian isolates. Similarly, sequences of avirulence genes in an international collection of 205 *L. maculans* isolates were recently published (Van de Wouw et al., 2023). Although powerful and exhaustive, such an approach cannot be performed on hundreds of isolates every year. In addition, this method necessitates first to isolate and purify each isolate.

Here, we developed a tool based on MiSeq sequencing technology, with two levels of multiplexing. First, eight *AvrLm* genes and one control gene (*Actin*) are amplified together in a multiplex PCR. Second, the multiplex-PCR is done on pools of phoma leaf spots, the typical *L. maculans* symptoms on *B. napus* leaves, to increase the size of the population assessed and to avoid the *L. maculans* isolation step. PCR products of pooled samples are sequenced using MiSeq technology and the proportions of each allele of each avirulence genes in the pooled sample are determined. The tool is named here “MPSeqM tool » (Multiplex on Pools and Sequencing with MiSeq). After the development and validation of the tool on DNA pools of known isolates, we applied the method to samples composed of pools of leaf discs centred on phoma leaf spots from experimental fields. The MPSeqM tool was applied to pools of 32 to 96 different leaf spots (possibly corresponding to as many different individuals, Gout et al., 2006) and the results were compared to phenotyping data of isolate populations from the same fields and the same sampling date (Balesdent et al., 2023). This approach not only allowed us to define a robust protocol for large-scale monitoring of virulence in the field, but also revealed a diversity of situations between sites, between *AvrLm* genes, and identified many unknown and rare alleles.

## MATERIALS AND METHODS

### Isolates

Seventy-one *L. maculans* isolates were chosen, including the reference isolate JN3 (v23.1.3), based on their polymorphism in their avirulence genes (Supplementary Tables S1 and S2; Gautier et al., 2023). The isolate X83.51 was generated following an *in vitro*-cross aiming at accumulating Avr gene deletion in a single isolate. *AvrLm1*, *AvrLm4-7*, *AvrLm11* and *AvrLmS-Lep2* are deleted in this isolate. X83.51 was used as a negative control for these genes and in serial DNA dilutions with variable proportions of deleted alleles.

### Field samples

The oilseed rape variety Westar, devoid of resistance genes, was sown at seven sites in autumn 2019 and at one site in 2020 (Table 1), with two to five plots per site. Individual leaves with typical *L. maculans* leaf spots were collected in autumn (300 per site, all collected from distinct plants). Part of these leaves was previously used to isolate around 80 single-pycnidium isolates per site for pathotyping on a differential plant set (Balesdent et al., 2023). In parallel, leaf disks (one disk per leaf) centred on a single leaf spot were cut using a cork-borer (1 cm diameter). Leaf disks were stored in 24-weel microplates at -20°C until further use (Supplementary Figure S1). Before DNA extraction, leaf spots from a given site were pooled by 24 or 32 in one Falcon tube and freeze-dried for 24 hours. Three to six pools per site were prepared (Table 1).

**Table 1.**
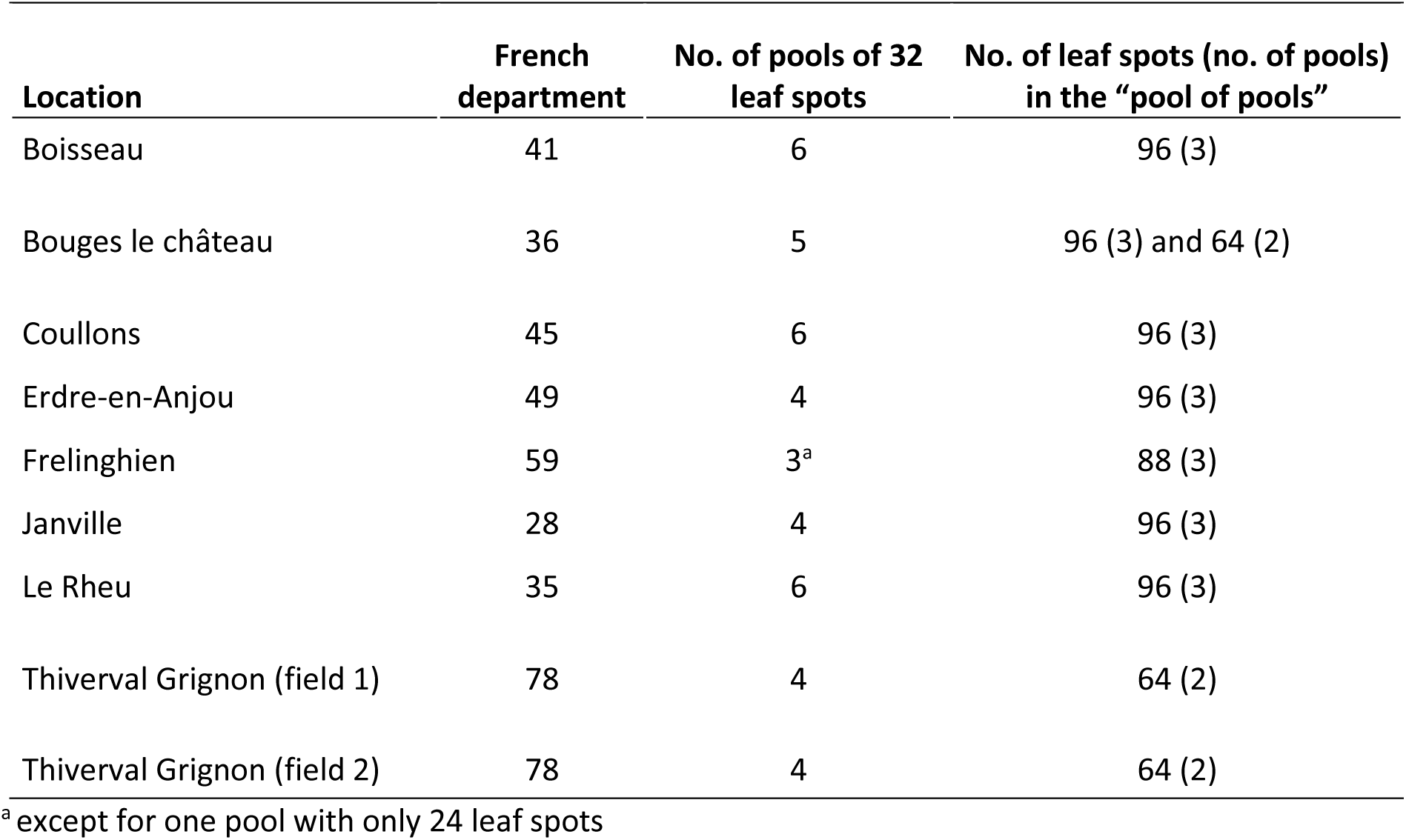
Characteristics of the sites sampled in autumn 2019 and 2020, and number of samples analysed per site.

| **Location** | **French**<br>**department** | **No. of pools of 32**<br>**leaf spots** | **No. of leaf spots (no. of pools)**<br>**in the “pool of pools”** |
| --- | --- | --- | --- |
| Boisseau | 41 | 6 | 96 (3) |
| Bouges le château | 36 | 5 | 96 (3) and 64 (2) |
| Coullons | 45 | 6 | 96 (3) |
| Erdre-en-Anjou | 49 | 4 | 96 (3) |
| Frelinghien | 59 | 3<sup>a</sup> | 88 (3) |
| Janville | 28 | 4 | 96 (3) |
| Le Rheu | 35 | 6 | 96 (3) |
| Thiverval Grignon (field 1) | 78 | 4 | 64 (2) |
| Thiverval Grignon (field 2) | 78 | 4 | 64 (2) |

<sup>a</sup> except for one pool with only 24 leaf spots

### Plant inoculation for preparation of DNA control

A selection of 32 *L. maculans* isolates with known avirulence alleles was inoculated on cotyledons of *B. napus* cv. Westar using standard protocols (Gautier et al., 2023; Supplementary Tables S1 and S2). Fourteen days after inoculation, leaf spots that developed on cotyledons were cut as described above. One spot per isolate was placed in a 50 mL Falcon tube, to constitute a pool of 32 leaf spots of known *AvrLm* allele composition. The pool was freeze-dried during 24 hours before DNA extraction.

### DNA extraction and manipulation

For each of the 71 *L. maculans* isolates, fungal DNA was extracted independently from 1 mL of crude conidial suspensions (between 10^8^ and 10^9^ spores mL^-1^) using the DNeasy 96 Plant Kit (Qiagen) following the manufacturer’s instructions (Gautier et al., 2023). Freeze-dried pools of leaf spots, whether from inoculations or from the fields, were crushed with a tungsten carbide bead (3 mm, Qiagen) on a Retsch MM400 mixer mill for 60 seconds at 30 Hz and DNA was extracted using the DNeasy Plant Mini kit (Qiagen). DNA was eluted in 100 µL of elution buffer AE (Qiagen). DNA quality was assessed with NanoDrop™ One (Thermo Scientific™) and DNA concentration was measured with Qubit™ 2.0 fluorimeter (Invitrogen™). After extraction, DNA of pools of 32 field leaf spots from the same field were mixed equimolarly to constitute “pool of pools” composed of mainly 96 leaf spots (Table 1).

### Control DNA samples

Four different controls were included, in triplicate, in each sequencing run: (i), multiplex PCR of JN3 DNA (termed JN3_DNA) used as a positive control; (ii), multiplex PCR of the pool of 32 leaf spots from inoculations with known isolates (termed Mix_LSpot); (iii), multiplex PCR of a pool of 71 *L. maculans* DNA (termed Mix_LM); and (iv), multiplex PCR of the 11 points of a dilution series for deletion quantification (termed C_dil_ser). The Mix_LM was prepared by mixing 50ng of DNA of each isolate and used to test the ability to detect all allelic variants in a pool (Supplementary Table S2). The dilution series was prepared by mixing DNA of isolates JN3 and X83.51 in proportions varying from 0 % to 100% (with a 10% increment).

### Multiplex PCR and MiSeq sequencing

Two technical criteria governed primer design, due to MiSeq technology and multiplexing; (i) the amplified region must be smaller than 460 bp for correct paired-end subsequent analysis; (ii) all amplified sequences in the multiplex PCR must be of similar length to avoid over-representation of the shortest sequences (maximum of 100 bp difference). Primers fitting these criteria were designed using the Primer3 website https://primer3.ut.ee/ (Table 2). The PCR reaction contained 40 ng of DNA, 1X Master Mix of Type-it Microsatellite PCR Kit (Qiagen), 4 μL Q-Solution, and 0.25 μM of each primer (except *AvrLm2* primers at 0.2 µM and *AvrLmS-Lep2* primers at 0.4 µM) in a total volume of 40 μL. PCR reactions were performed using an Applied Biosystems 9700 thermocycler. The amplification parameters consisted of an initial denaturation step at 95°C for 5 min; then 35 cycles of denaturation at 95°C for 60 s, annealing at 60°C for 90 s, and elongation at 72°C for 60 s; and a final extension step at 72°C for 10 min. Three independent PCRs were performed for each sample to test the repeatability and robustness of the method. The PCRs were verified on 3% agarose gels and sent to the GenoToul platform (INRAE Toulouse) for Illumina MiSeq sequencing using the chemistry V3 (2x250pb).

**Table 2.**
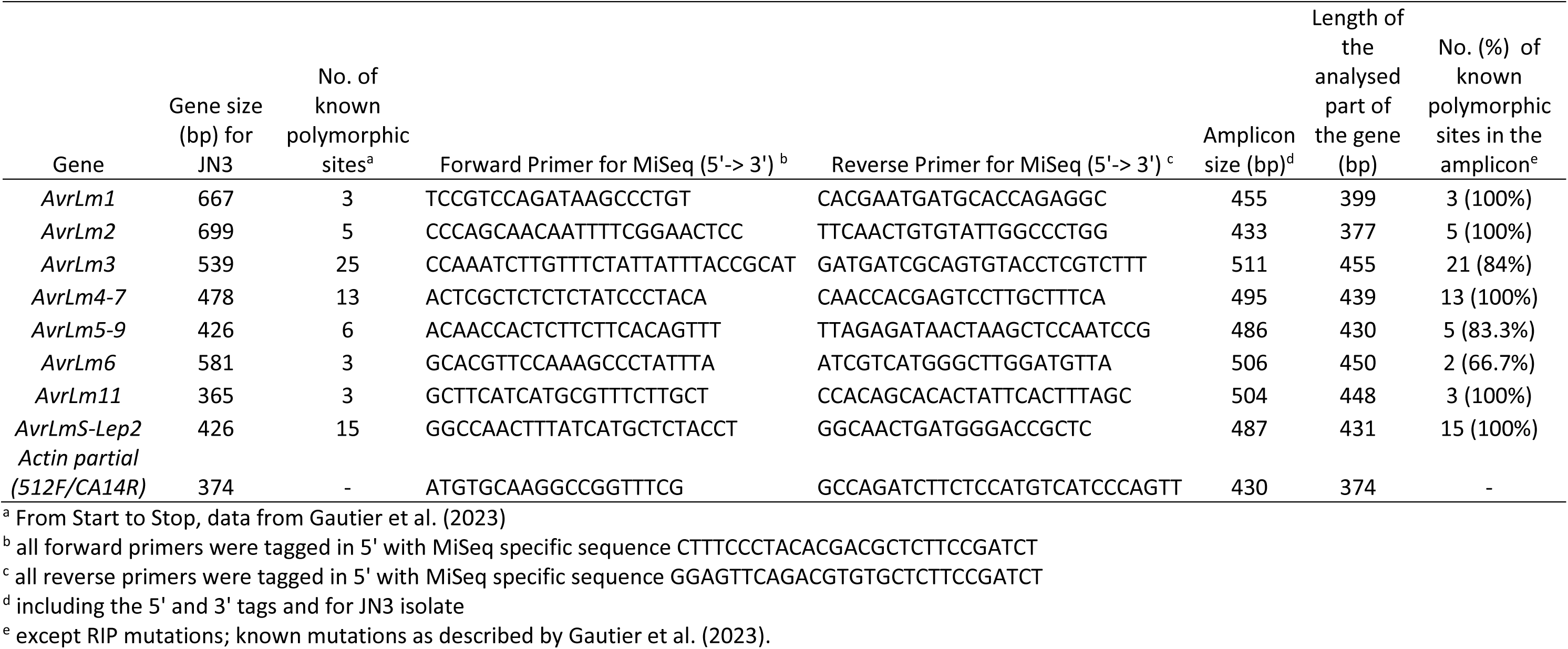
Characteristics of the eight *AvrLm* and *Actin* gene regions amplified with MiSeq primers.

### AvrLm allele database

At the start of the project, the *L. maculans AvrLm* database included the 87 alleles and 63 isoforms identified following the sequencing of 11 avirulence genes in a collection of French and reference isolates (Gautier et al., 2023) and available at https://bioinfo.bioger.inrae.fr/portal/data-browser/public/leptosphaeria/avirulence_genes. Additional *AvrLm3* and *AvrLm4-7* alleles described in Balesdent et al. (2022) were added to the database.

### Analysis of sequence dataset: pairing, filtering, processing and assigning an allelic classification to sequences

First, for each sample, forward (R1) and reverse (R2) reads were paired using the tool Pear - Paired-End read merger (Galaxy Version 0.9.6.2) of Galaxy pipeline (https://vm-galaxy-prod.toulouse.inrae.fr/galaxy_main). Then, the data were processed gene by gene. For each gene, only sequences with the expected primer sequences were retained. A stringent pipeline (Boutigny et al., 2019) was used to filter and process these paired sequences, utilising MOTHUR v.1.36.1 (Schloss et al., 2009) and parameters adapted to each gene (Supplementary Text T1). Sequences with low quality (shorter or including low-quality scores) were removed (Supplementary Text T1). After preprocessing the sequences, sequence files were merged, and unique sequences represented by singletons or 2 reads were removed. Chimeric sequences were removed with the Uchime tool (Edgar et al., 2011) available in MOTHUR, using self as the reference. Uncorrected pairwise distances were determined between the remaining sequences with the Needleman alignment method (Needleman et al., 1970) with match 1.0, mismatch -1.0, gapopen -2.0 and gapextend 1.0 using 1000 iterations. The distance matrix was then clustered with the average neighbor algorithm to assign sequences to operational taxonomic units (OTUs). Using MOTHUR’s classifier (Wang et al., 2007), the representative sequence for each OTU was assigned to the allele of the corresponding *AvrLm* gene from the custom *Lmb_avrlm_act genes* reference database using no divergence. An OTU count table was obtained containing OTU number, read count per OTU per sample, representative sequence name, representative sequence, and OTU representative assignation (Figure 1). In a final step, the Galaxy blastn tool was used, to compare the OTU sequences retained against *Lmb_avrlm_act genes* references database and distinguish the known alleles with 100% identity in length and sequence to new alleles with less than 100% identity in sequence.

**Figure 1.**
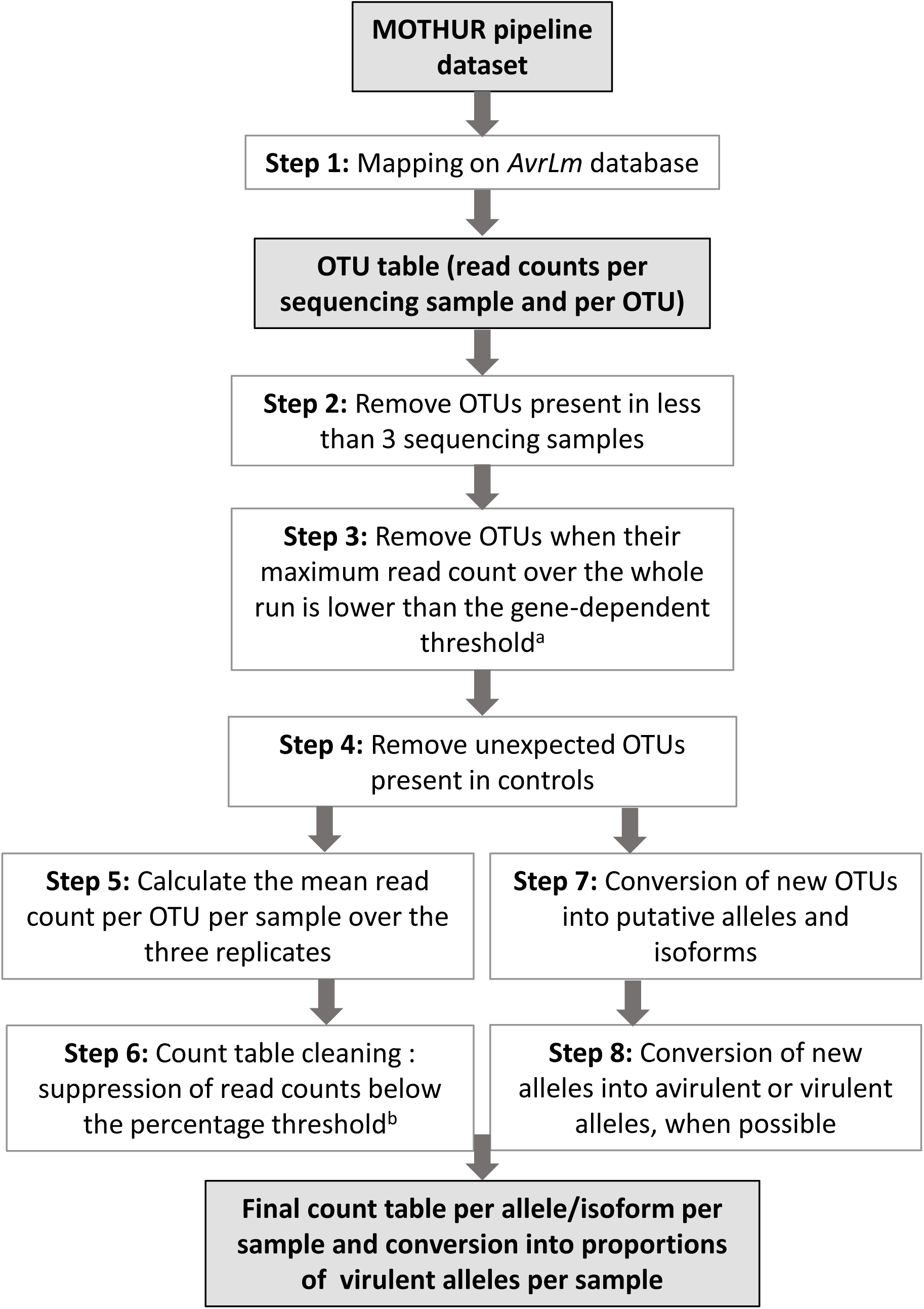
Schematic representation of sequence analyses and dataset cleaning. Grey boxes correspond to output files at specific stages of the analysis. ^a^, this threshold (“maximum read number threshold”) removes background OTUs due to sequencing errors and is based on unexpected values observed for control DNAs; ^b^, this threshold (“percentage threshold”) is based on the relative proportion of an OTU in an field sample and its value is based on relative proportions observed in control DNAs.

### Strategy for the selection of relevant OTUs and database cleaning

After the mapping step, a very high number of OTUs were identified (Table 3), most of which correspond to artefacts due to PCR or the MiSeq technology. Three filters were sequentially applied (Figure 1) to keep only reliable OTUs as follows: (i), keep only OTUs present in at least three samples, due to the use of triplicates for each DNA sample; (ii), remove all OTUs for which, whatever the sample analysed, the read numbers were below a threshold. This threshold is named “Maximum read number threshold” and is gene-dependent (Table 3) (iii), remove OTUs that were detected in controls but should not be present. After this OTU selection step, we calculated the mean read number of the three replicates for each OTU and DNA sample. Finally, the resulting count table was simplified by considering an OTU absent in one sample if its proportion in this sample (i.e., read count of the OTU divided by total read counts for all OTUs for this gene) was lower than a threshold (“proportion” threshold), specific for each gene. This percentage threshold is determined based on the expected and observed OTU proportions in controls (Table 3, Figure 1).

**Table 3.**
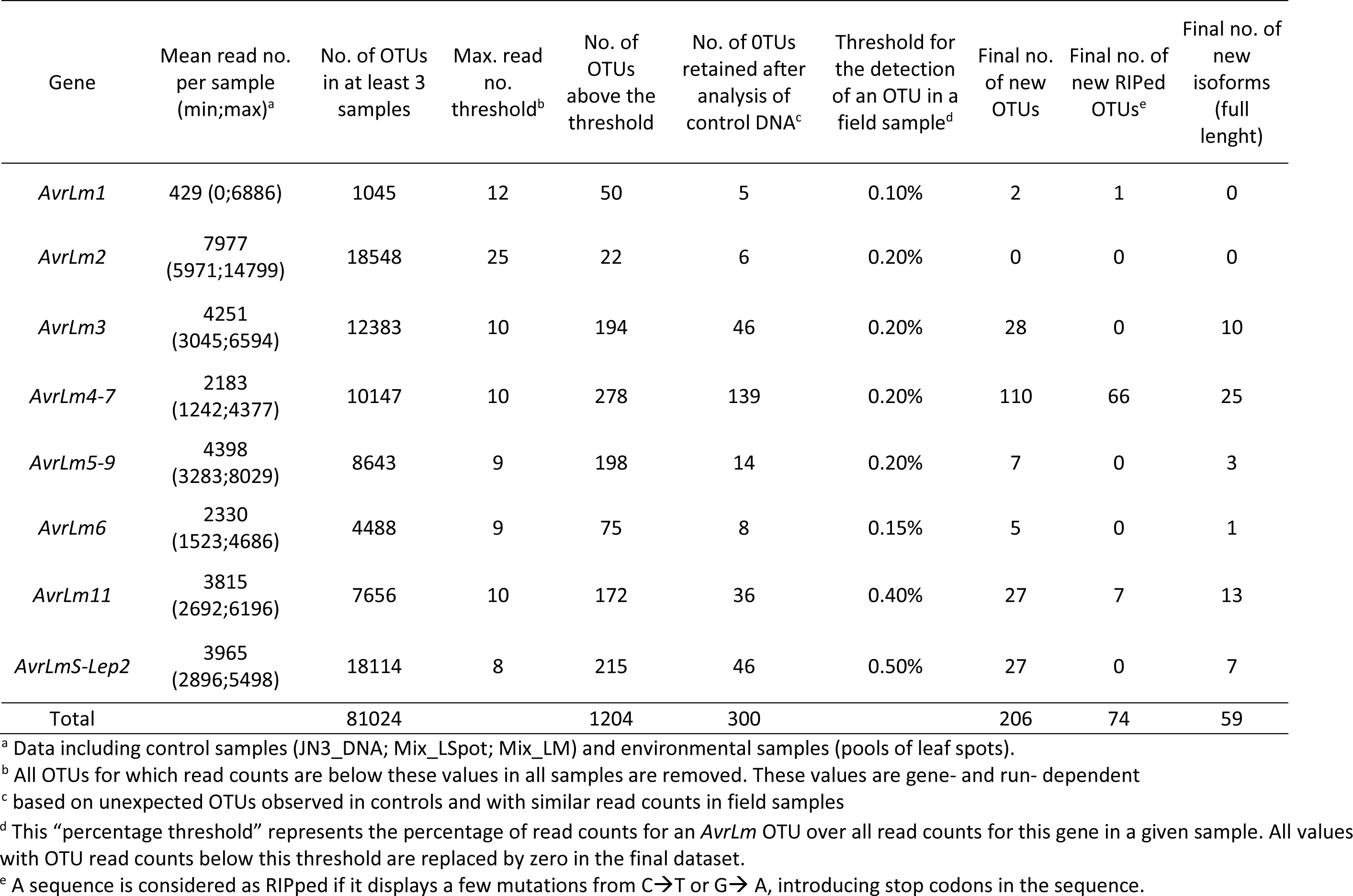
Sequencing statistics including numbers of OTUs retained at each filtering step.

### Analysis of new AvrLm alleles

When a new OTU was detected, the missing parts of the gene sequence, if any, (i.e., when primers were within the genes) were completed by the corresponding 5’ or 3’ sequences of the JN3 reference isolate to generate a new putative allele. Introns were removed and the sequence was translated with the Expasy translate tool (https://web.expasy.org/translate/) to generate an extrapolated, putative new isoform (Figure 1).

### Validation of new alleles

DNA of one leaf spot pool from the Coullons site (Table 1) displaying a high proportion of new OTUs was used to clone and validate putative new alleles. After *AvrLm4-7* PCR amplification with external primers (Gautier et al., 2023) the PCR products were cloned into pGEM**®**-T Easy Vector System (PROMEGA) using standard protocols. 101 colonies were recovered. The *AvrLm4-7* insert was re-amplified in colonies with the plasmid standard primers SP6/T7 and sequenced by Eurofins Genomics (EUROFINS, Ebersberg, Germany) using SP6 external primer.

## RESULTS

### Primer design and reproducibility of multiplex PCR and sequencing

We designed a set of primers fitting technical constraints and amplifying parts of *AvrLm* genes containing a maximum of known polymorphic sites and all codons encoding a cysteine (Table 2). These primers amplified part of the genes containing between 66% and 100 % of the known mutations (Table 2, Supplementary Figure S2). A few known variants were not detected, with no or low consequences for the diagnosis of the allele as an avirulent or a virulent one. For instance, four *AvrLm3* variants cannot be detected as mutations are outside the amplified region (Table 2). However, these mutations are either in the last intron or in the peptide signal-coding region, without impact on the mature protein (Supplementary Fig S2-C).

Using these primers, we first tested the MPSeqM tool with the concomitant amplification of four, five, or nine genes in a single PCR. Sequencing results were similar whatever the number of genes amplified together (data not shown) and we thus choose the nonaplex amplification in all further experiments. MPSeqM of control DNAs (JN3_DNA, Mix_LSpot, and Mix_LM) revealed an efficient and reproducible amplification level of all genes (Figure 2 and data not shown). The less amplified gene was *AvrLm6*, followed by *AvrLmS*, still representing more than 1250 reads each in each control replicate.

**Figure 2.** Efficiency and reproducibility of MiSeq sequencing following multiplex PCR amplification of *Leptosphaeria maculans* DNA controls. A, the JN3 reference, single isolate, DNA; B, pooled leaf symptoms from 32 known isolates. Replicates (REP) correspond to independent PCR experiments. Values are percentages of reads matching each amplified gene.

### Detection and quantification of deletions

To estimate the proportion of deleted alleles of a given *AvrLm* gene in a field sample, we used serial dilutions of the DNA of an isolate deleted for this gene with the DNA of an isolate with the gene present. The theoretical proportions of the *AvrLm* gene in the dilution series were compared with the estimated proportions calculated as the total number of reads mapping to this gene over the total number of reads mapping to the *L. maculans Actin* gene, which is a single copy and invariant gene. The two values were proportional and allowed us to define standard curves of variable shapes depending on the gene (Figure 3). We approximated the curves to a polynomial equation, whose parameters were used to calculate the proportion of undeleted events and, by difference, the deleted ones, in a pooled sample. For instance, the expected proportion of deleted *AvrLm1* in the Mix_LSpot is 87.5% ([exact confidence interval [71-96]) and the estimated proportion of the deletion calculated with the standard curve equation is 84.5% (Supplementary Table S3).

**Figure 3.**
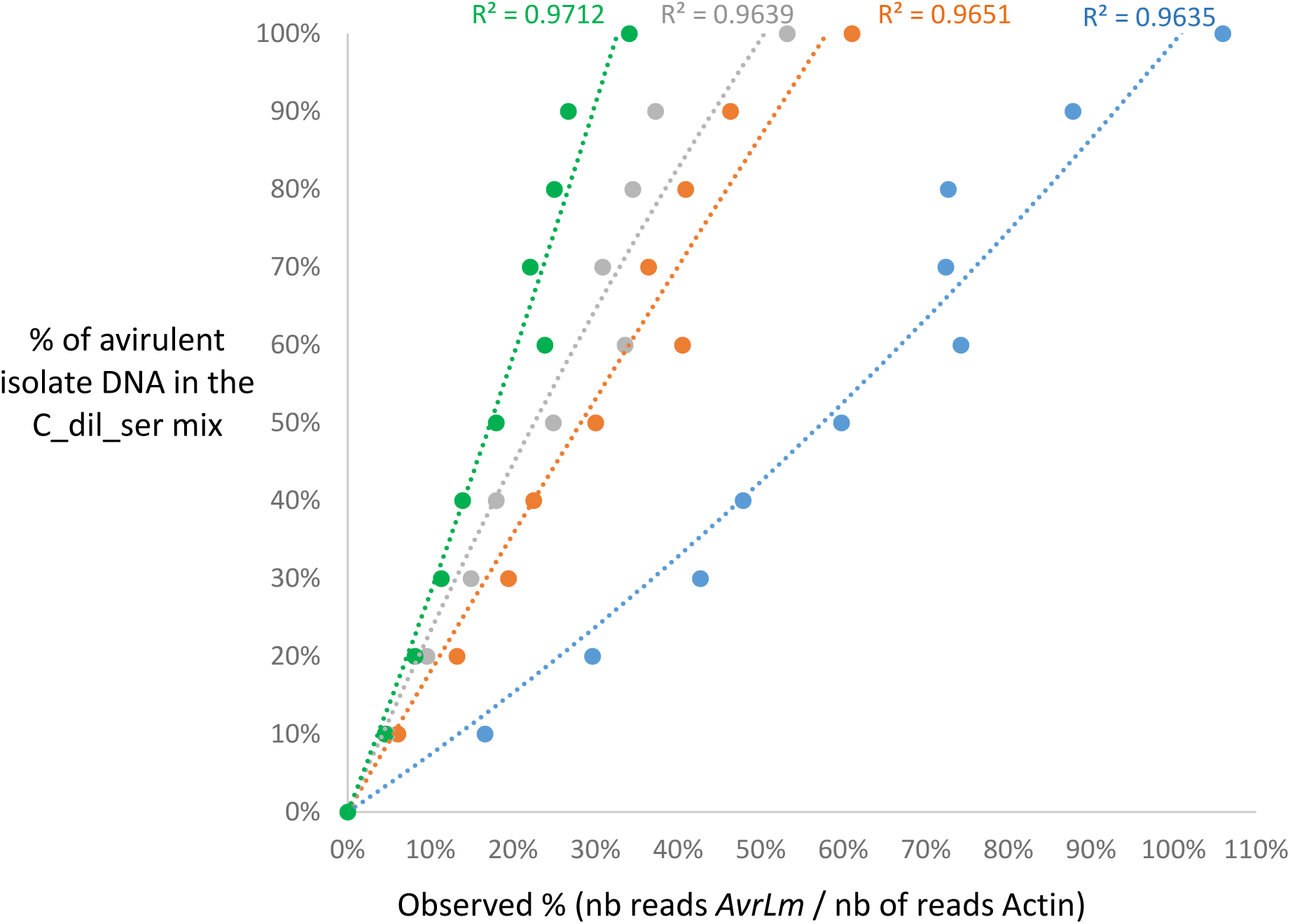
Serial dilution curves for *AvrLm* deletion quantification. A mix of DNA of two isolates differing by the presence or absence of four avirulence genes created deletion proportions ranging from 0% to 100%. The observed proportion of the gene in the mix was estimated with the ratio between the read numbers for this gene in the sample over the *Actin* read numbers. In blue, *AvrLm1*; orange, *AvrLm4-7*; grey, *AvrLm11*; green, *AvrLmS-Lep2*. Polynomial regression curves and R^2^ are displayed for each gene.

### Threshold determination for OTUs assignation

The use of triplicates for each sample and four types of controls of known allele composition was essential to determine the thresholds used to keep only reliable OTUs and suppress potential PCR or sequencing artefacts. Mainly, the “Maximum number of reads” threshold allowed us to reduce drastically the number of OTUs (1204 OTUs in total, starting from 81024 OTUs kept with the criteria of presence in at least three sequencing samples, Table 3). The controls with the reference JN3 isolate or with DNA mixes of known isolates also allowed us to remove all OTUs still unexpectedly found in these controls, which represented 3/4 of the remaining OTUs. Altogether, 300 reliable OTUs (all genes) were kept, among which 206 were not in the database, 74 of which corresponded to highly mutated versions of the genes by RIP (Table 3). Lastly, depending on the gene, OTUs representing less than 0.1 to 0.5% of the total read counts for this gene in a field sample (“proportion” threshold) were considered absent from this sample (Table 3).

### Validation of the method on control samples

The previous steps allowed us to select reliable OTUs and to determine the proportion of deleted alleles for *AvrLm1*, *AvrLm11*, and *AvrLm4-7*. Using this cleaned dataset, we compared the proportion of alleles (OTUs) detected in control samples to the expected ones (Figure 4, Supplementary Table S3). In the Mix_Lm control, including 71 isolates, 20 alleles out of 90 were not detected, but 19 of them corresponded to a rare occurrence (only one isolate among 71). In addition, most of them (13) were RIPped alleles. In the Mix_LSpot control, only 3 out of 47 expected alleles were not detected, all corresponding to RIPped alleles for *AvrLm1* or *AvrLm4-7*. All other alleles, including those present in only one isolate out of 32, were detected (Supplementary Table S3). Overall, observed and expected proportions correlated very well (R^2^>0.98). Only one *AvrLm4-7* allele (the deleted allele) was overestimated in the Mix_LM control, possibly due to the lack of amplification of some RIPped alleles. Apart from this allele, all estimated values were within the exact confidence interval determined for sample sizes of 32 (Mix_LSpot) or 71 (Mix_LM) individuals (Figure 4).

**Figure 4.**
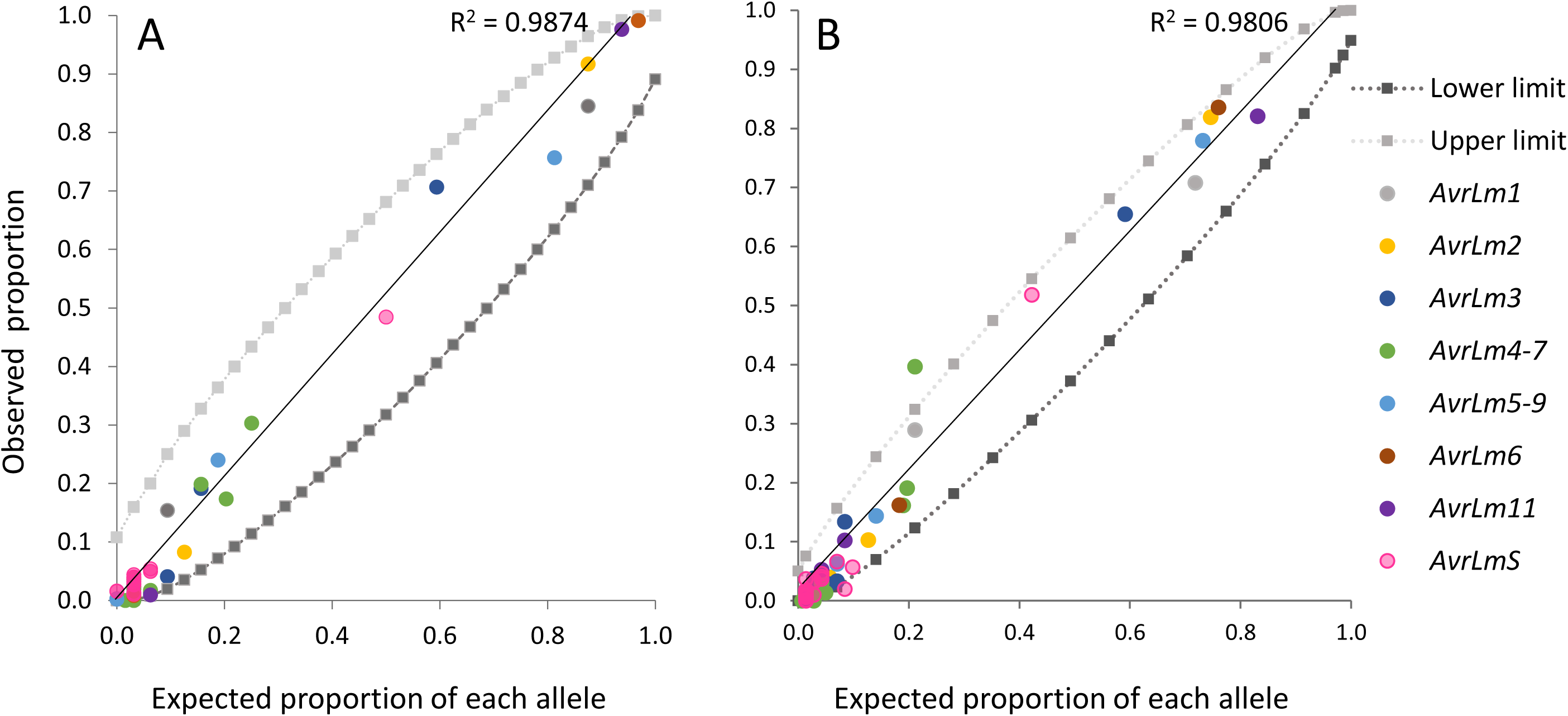
Expected and observed proportions of alleles of eight *L. maculans* avirulence genes in DNA pools. A, DNA of a pool of 32 leaf spots from known isolates and B, pool of DNA of 71 known isolates. The eight *AvrLm* genes were amplified in multiplex PCR with three independent PCR replicates, and then sequenced with MiSeq. Each dot correspond to the mean proportion in the pools of each allele (OTU) over all alleles of every *AvrLm* gene, including deleted alleles, and the colour code on the left refers to the gene. The lower and upper limits correspond to the exact confidence interval around the expected value for a mix of 32 (A) or 71 (B) individuals. Detailed values are in Supplementary Table S3.

### Analysis of unknown field samples: identification of new alleles

The MPSeqM tool, validated on controls, was then used on 42 field samples of pooled leaf spots corresponding to 1337 *L. maculans* symptoms from eight French regions (Table 1). This analysis first revealed several still undescribed alleles for most of the *AvrLm* genes studied.

Such new alleles were considered as real alleles, not sequencing artefacts, when they combined the following characteristics: (i) they were detected in some samples only, not as a background signal in all samples including controls; (ii) they were consistently detected in the three independent technical replicates of one given field sample; (iii) their read numbers were higher than the gene-dependent threshold value determined for each gene; and (iv) they were detected both in one “pool of 32 leaf spots” and in the corresponding “pool of pools”, when available, with consistent proportions (Figure 5). Using these criteria, a total of 206 new alleles were detected.

**Figure 5.**
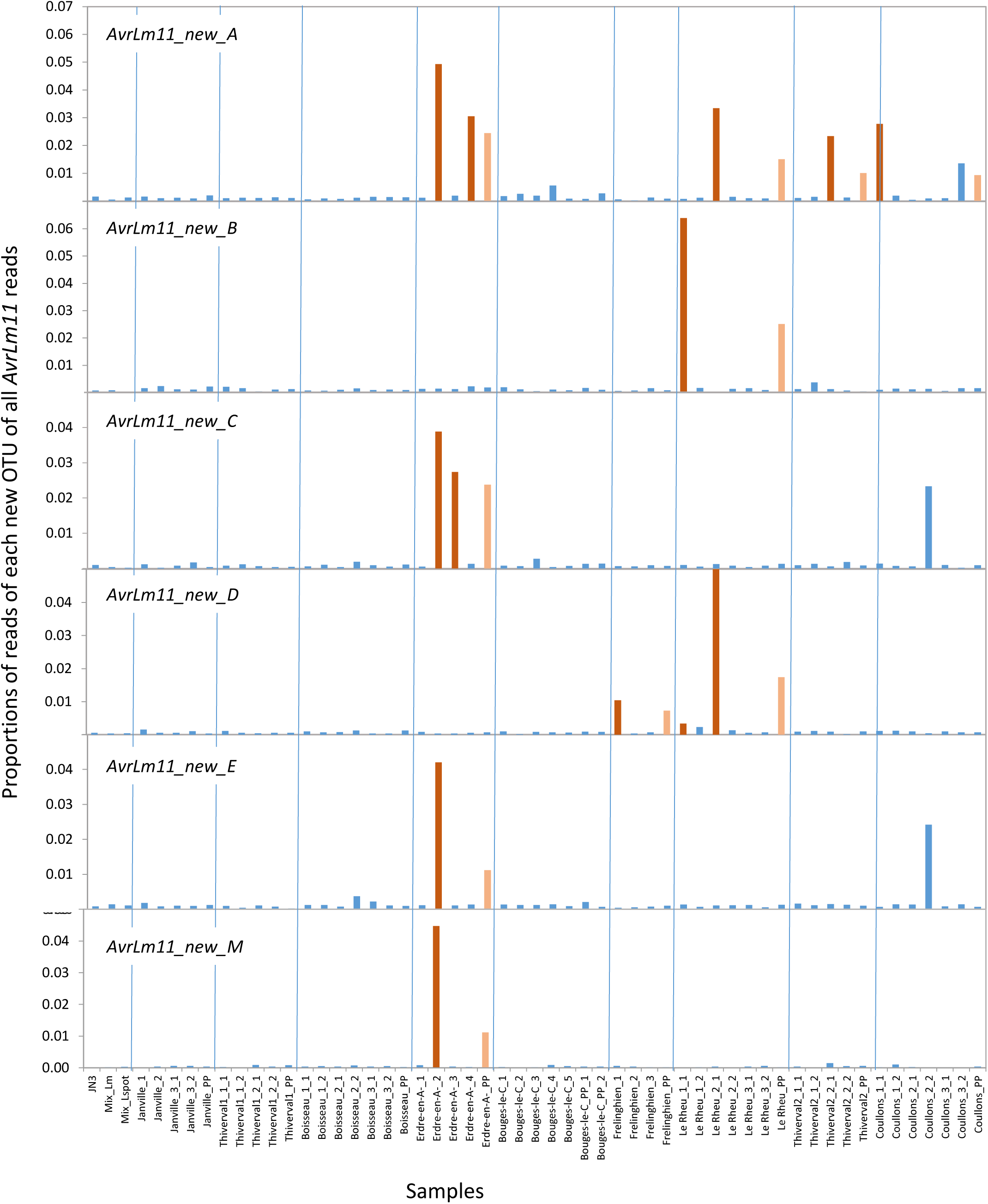
An example of detection of new alleles. Percentages of new OTUs over all *AvrLm11* OTUs are given for the three control samples (left) and the 52 field samples from nine sites, separated by the vertical blue lines. Bars in dark orange are those of pools of 32 leaf spots that are also in the “pools of pools “of 96 leaf spots (pale orange).

### Analysis of unknown field samples: gene-by-gene analysis

#### AvrLm1

Only one out of the five previously described *AvrLm1* alleles was detected in field samples, allele *AvrLm1_0_A.* In addition, we detected two new alleles, one RIPped allele with 14 substitutions and one allele with a stop codon in the middle of the sequence. These two new alleles were considered as virulent, as were deleted isolates whose proportions were estimated using the deletion curve. Altogether, the MPSeqM analysis of field samples correlated well with pathotyping; 1.8% of the population was found avirulent on *Rlm1* by pathotyping (Balesdent et al., 2023), while the MPSeqM analysis of populations from the same sites detected 2.1% of such isolates. With both approaches, the site with the highest proportion of avirulent isolates was that of Le Rheu (Supplementary Figure S3). With the MPSeqM tool, it was possible to reproducibly detect low percentages of avirulent isolates in all sites, while, by phenotyping 30 isolates per site, such isolates were not detected (Balesdent et al., 2023).

#### AvrLm2

No new *AvrLm2* allele was detected in field samples. The *AvrLm2_0_A* was largely prevalent, with proportions ranging from 91% to 100% (average across all pools = 97.3%, Supplementary Figure S4) depending on the pool. The *AvrLm2_1_A* allele was found in all sites at a low frequency (0 to 8% per pool, average across all pools = 2.2%). *AvrLm2_2_A* was found in six sites (10 pools out of 42; between 0 and 6.9%; average across all pools = 0.52%). The other known alleles, including all avirulent ones, were never detected in the pools.

#### AvrLm3

From the 24 previously described *AvrLm3* alleles, 13 were detected in the field samples. Twenty-eight new OTUs were also detected, being either rare (present in one sample only) or more frequent (present in up to 23 samples), allowing us to identify in total 41 different OTUs for *AvrLm3*. The new alleles could represent up to 8.5% of the OTUs in a given sample. These new alleles encoded for ten putative new isoforms of the AvrLm3 protein. Based on their alignment with previously described AvrLm3 isoforms, we inferred that six among them potentially correspond to avirulent isoforms and four to virulent ones (Supplementary Figure S5). Despite the allelic diversity detected in field samples, four alleles represented 96% of the population, with allele *AvrLm3_A1* being by far the most abundant (80.2%), followed by *AvrLm3_C* (11.8%) (Supplementary Figure S6), consistent with previous surveys and sequencing (Gautier et al., 2023).

#### AvrLm4-7

MPSeqM results for field samples displayed 110 new *AvrLm4-7* alleles (Table 3). Most of them (66) corresponded to RIPped alleles with mutations introducing stop codons in the sequence and thus corresponding to virulent alleles, also unmasking *AvrLm3*. Seven sequences displayed one or two bp deletions, also encoding for truncated (virulent) proteins. A few (17) were new alleles of known (virulent or avirulent) isoforms, however, 25 new isoforms were detected. The amino acid (AA) sequence of these new isoforms was compared to previously described ones (Balesdent et al., 2023) and the corresponding phenotypes were more or less easily deduced depending on the AA substitutions. One isoform putatively corresponded to an A4A7 (conferring avirulence on both *Rlm4* and *Rlm7*) isoform, six alleles for a3a7 isoforms (conferring virulence on *Rlm4* and *Rlm7* and on *Rlm3* by masking effect), four for a3A7 (avirulent on *Rlm7* only) isoforms. For the remaining 14 isoforms, it remained impossible to conclude.

#### AvrLm5-9

Due to primer design, alleles *AvrLm5-9_0_A* and *AvrLm5-9_0_C* (encoding for the same isoform) cannot be distinguished. In field samples, seven new variants were detected. Only three of them corresponded to putative new isoforms of the protein; one displayed one AA change in the signal peptide compared to the AvrLm5-9_0 isoform without any modification of its secretory properties (data not shown). The two others displayed only one AA change each (R63L and K101Q, respectively), whose impact on the phenotype cannot be determined. Among the two alleles encoding avirulent isoforms on *Rlm9*, isoform *AvrLm5-9_4* was never detected in any of the 42 pools, and *AvrLm5-9_1* was found in only nine pools with very low proportions (<0.27%, close to the 0.2% detection limit, 0.06% on average over all pools), confirming the near-fixation of the virulence phenotype toward *Rlm9* in French *L. maculans* populations (Balesdent et al., 2023 Gautier et al., 2023). Similarly, alleles with stop codons in the coding sequence were never detected. Depending on the site, from two to six alleles were detected (Supplementary Figure S7).

#### AvrLm6

Field samples displayed five new OTUs. They were however rare, present in one or two sites only with proportions ranging from 0.6% to 4.6% (Supplementary Figure S8). Each new OTU displayed only one mutation. One was in an intron; three were silent and only one of them introduced a G47C substitution, and thus a new isoform. The impact of this mutation on the phenotype cannot be determined yet. Allele *AvrLm6_0_A* was largely prevalent, ranging from 91.7% to 100% depending on the sample, followed by allele *AvrLm6_1_A* or *AvrLm6_3_A*, two alleles that cannot be discriminated with the current primers (between 0% and 4.1%).

#### AvrLm11

For *AvrLm11*, two new OTUS displayed a shorter-than-expected size (one or two bp). These gaps were either in the intron or after the stop, thus without impact on the isoform of the protein. In addition, 27 new OTUs were detected in at least one field sample, displaying from one to seven mutations along the gene. These new OTUs were often specific to one sampling site (Figure 5) and, depending on the site, from one to 14 new OTUs were present. Most of the mutations (72%) in these OTUs were typical of RIP mutations (C->T and G-> A). In these OTUs however, the number of mutations was limited (from 1 to 7 mutations per *AvrLm11* OTU) compared to what is seen in RIPped alleles for other genes like *AvrLm4-7* (from 1 to 44 mutations, mean=20.2, data not shown). Despite this low number of mutations per allele, these RIP mutations introduced stop codons in seven OTUs at positions 29 or 74 of the protein leading to truncated versions of the protein that thus corresponded to virulent alleles. For 11 OTUs, the mutations were silent, including one mutation in the intron or three mutations after the stop codon, and corresponded to the avirulent AvrLm11_0 isoform. The remaining OTUs corresponded to 13 new isoforms. One of them displayed a mutation in a codon encoding a cysteine, therefore generating a de-structured protein and the probable loss of avirulence. For the other 12 new isoforms, we could not determine yet the impact of these mutations on the interaction phenotype. However, they represented only 0.7% of the detected alleles in average, but could reach up to 7.8% in pools displaying numerous new alleles. Altogether, alleles encoding the AvrLm11_0 isoform were largely prevalent in the population (94.6% in average), while isoforms AvrLm11_1 or AvrLm11_2, previously described in Canada, USA or New Zealand (Chen et al., 2020; Van de Wouw et al., 2023) were never detected. *AvrLm11* deletion was estimated at 3.1% in average, ranging from 0% to 15.4% depending on the sample.

#### AvrLmS

As already observed following the sequencing of individual isolates (Gautier et al., 2023; Van de Wouw et al., 2023), a high level of polymorphism was detected for *AvrLmS*. Forty-one alleles were detected here, with 27 new alleles compared to previous data (Gautier et al., 2023). The proportions of each *AvrLmS* allele greatly varied between samples including within one given site (Supplementary Figure S9). Two new alleles displayed one mutation introducing a stop codon in the sequence, as previously detected in virulent isolates (Gorse M. and Gautier A., unpublished data; Van de Wouw et al., 2023). These virulent alleles were detected in only one pool from the Boisseau site at low frequency. Nineteen new alleles displayed silent mutations and encoded avirulent isoforms. Three alleles displayed non-synonymous mutations leading to AA substitutions in the signal peptide without modification of the mature (avirulent) protein. Only three alleles displayed non-synonymous mutations (R41Q, S49L and E71G) for which the impact on the phenotype could not be determined. They represented in average 0.56% of the detected alleles (data not shown).

### Analysis of unknown field samples: comparison between 32 or 96 leaf spots in a pool

We compared the proportions of each detected allele in pools of 32 leaf spots or in “pools of pools” containing up to 96 leaf spots. As exemplified for *AvrLm3* for which many alleles were detected, the proportion of each allele estimated from both pool sizes were highly correlated (R^2^=0.9996, Supplementary Figure S10). However, when focusing on alleles present at low percentages in the pools (<2%) the correlation was lower (R^2^=0.859), due to OTUs that were not detected in pools of 96 leaf spots (detection limit) but also due to OTUs detected in pools of 96 but not in pools of 32, probably corresponding to PCR or sequencing artefacts (Supplementary Figure S10).

### Analysis of unknown field samples: comparison between genotyping and phenotyping for virulence

Using leaf samples from the same site and the same sampling date, we compared the percentages of avirulent isolates in *L. maculans* populations as estimated following inoculation tests on a plant differential set (Balesdent et al., 2023) or using the MPSeqM tool (this study). For most avirulence loci, the inference of the phenotype (virulence or avirulence) from the allele sequences or deletion proportions was possible (see above). As regards *AvrLm3*, the situation is more complex since the avirulence phenotype is masked whenever *AvrLm4-7* is present, at least for some *AvrLm4-7* alleles (Balesdent et al., 2023). To determine the proportion of avirulent isolates on *Rlm3*, we multiplied, for each pool, the percentage of *AvrLm3* avirulent alleles by the percentage of *AvrLm4-7* alleles unable to mask the *AvrLm3-Rlm3* interaction. Altogether, phenotypic or MPSeqM analyses gave very close estimates of the proportions of avirulent isolates for the interactions investigated, including the complex *AvrLm3-Rlm3* interaction (Figure 6A). For instance, site by site comparison of MPSeqM or phenotypic estimation of virulent isolates on *Rlm7* perfectly correlated (Supplementary Figure 11).

**Figure 6.**
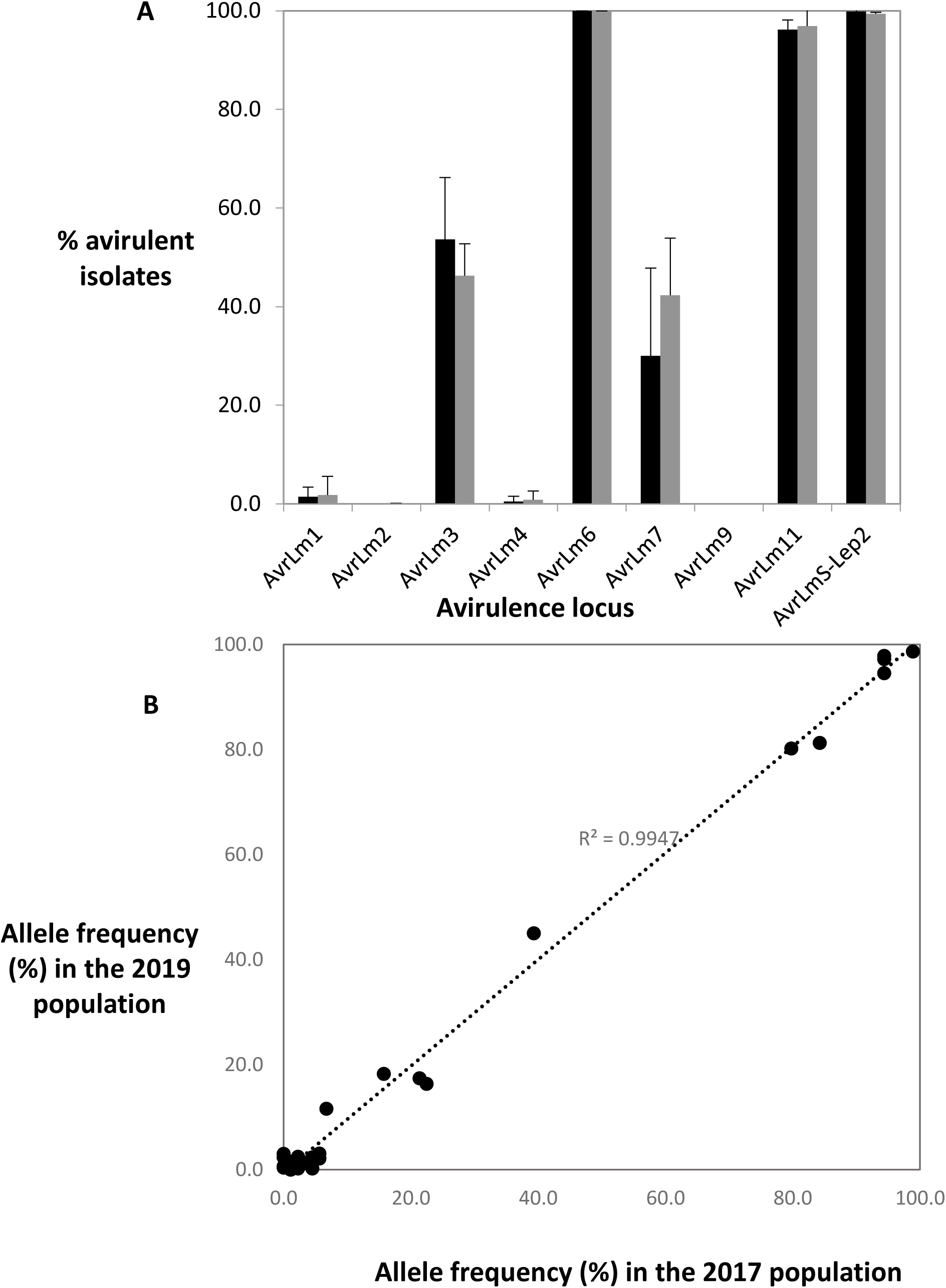
Validation of the MPSeqM tool on field samples. A, Comparison between percentages of avirulent isolates in French populations of *Leptosphaeria maculans,* estimated by phenotyping or genotyping. Black, data from phenotyping 690 isolates from 9 sites (Balesdent et al, 2023); grey, data from MultiplexPCR-Miseq analysis of 42 pools of leaf spots from the same sites (including a total of 1337 leaf spots). Values are means and standard errors over all sites. B, Comparison between allele frequencies (all genes) estimated by Sanger sequencing of individual isolates (2017 population, Gautier et al., 2013) and by the MPSeqM tool of pools of leaf spots (2019 population).

### Analysis of unknown field samples: comparison between Sanger sequencing and the MPSeqM tool

We previously estimated *AvrLm* allele frequencies in French *L. maculans* population by Sanger sequencing of full-length avirulence genes in a representative population of 89 isolates collected in 2017 (Gautier et al., 2023). These proportions were compared with those determined by the MPSeqM tool using averaged frequencies of alleles over all 42 field samples. Proportions of alleles that are undistinguishable with the MiSeq primers (Like *AvrLm3_A_01* and *AvrLm3_A_09*) were first merged, and similarly, proportions of RIPped and deleted alleles for *AvrLm1* were merged. Altogether, 44 distinct alleles were detected for the eight *AvrLm* genes in the 2017 population. The percentages of these alleles estimated with the MPSeqM tool in the 2019 population or by Sanger sequencing of avirulence genes in the 2017 populations perfectly correlated (R^2^=0.995, Figure 6B). The difference between the two estimates of a given allele was low (1.8% in average). For only two *AvrLm4-7* alleles, differences between the two analyses were higher than 5%, *AvrLm4-7_2_A* (an avirulent allele) being lower in 2019 (16.3%) compared to 2017 (22.5%) and the deleted allele being higher in 2019 (45.0%) compared to 2017 (39.3%; Figure 6B). Indeed, these differences may simply reflect the evolution toward virulence of French populations between 2017 and 2019 due to the *Rlm7* selection pressure in France (Balesdent et al., 2022).

### Analysis of unknown field samples with the MPSeqM tool: allelic diversity per site

Considering all genes, from 80 to 118 distinct OTUs were detected per site. Relative to the number of leaf spots per site analysed, the site showing the highest diversity (number of distinct OTUs) was that of Frelinghein (Supplementary Figure S12). Depending on the sites, from 40% (Frelinghein) to 53% of the detected OTUs were new OTUs, the site with highest proportion of new OTUs being that of Coullons, mainly due to the high number of new *AvrLm11* OTUS found at that site (Supplementary Figure S12).

### Validation of the strategy including detection of new alleles

One pool was used to validate new alleles found with the MPSeqM tool by cloning. In this pool, 13 distinct allele of *AvrLm4-7* were detected and kept after the filtering steps with the MPSeqM tool. When cloning and sequencing the PCR products after *AvrLm4-7* amplification with external primers, all but two of these 13 alleles were recovered. The proportion of clones with these alleles correlated the proportion of reads detected after MiSeq sequencing (data not shown) and these clones represented about 50% of the sequences cloned. Among these alleles, six corresponded to new alleles compared to previously published alleles, and were detected in one to seven independent clones. However, the remaining 50% of the clones each displayed unique alleles (i.e., each representing less than 1% of the amplified sequences) that were either considered not present in the pool, because below the threshold (13 sequences), or even not detected in the dataset whatever the pool (33 sequences).

## DISCUSSION

In this paper, we describe the design, validation, and use of a molecular tool, termed MPSeqM, for large-scale analysis of polymorphism of eight *AvrLm* genes of *L. maculans* intending to determine proportions of virulent isolates toward the corresponding *B. napus Rlm* genes in field samples. It combines four steps: bulk DNA extraction of *L. maculans* symptoms from the field; a multiplex-PCR step amplifying eight *AvrLm* genes and *Actin* as an invariant gene; illumina sequencing of the PCR products; and lastly, blasting reads on an *AvrLm* variant database and estimation of proportions of each variant in the bulks. The tool was first evaluated on DNA pools of known samples and was found to be versatile, reproducible, quantitative, and efficient for the detection of rare and new allelic variants.

As regards versatility, the possibility to characterize simultaneously from four to eight *AvrLm* genes with identical results opens the possibility to modify the set of genes analysed depending on the characteristics of the populations to survey. For instance in France, *AvrLm2* is nearly invariant with fixed virulent alleles, rendering *Rlm2* useless (Balesdent et al., 2023, Gautier et al., 2023). The MPSeqM tool could thus be adapted by replacing *AvrLm2* amplification with other genes of interest, like *AvrLm10* or *AvrLm14*. Alternatively, fewer genes could be analysed simultaneously, to reach deeper sequencing per gene or to analyse more samples for the same sequencing cost.

Sequencing results were very reproducible both qualitatively and quantitatively between the three replicates within one sequencing run. Independent runs also gave identical estimates of proportions of each allele (data not shown). The quantitative dimension of the tool was evident, not only when comparing the MPSeqM results with expected proportions of each alleles in control bulks of known composition, but also when comparing MPSeqM results from bulks of field samples with previously established data, whatever the criteria compared (proportion of virulent isolates per site, or proportions of allelic variants per gene).

Three main questions had to be addressed to validate the tool: (i) what could be the best sample size (number of leaf spots) within each pool for field surveys, (ii) whether gene deletions could be quantified, and (iii) whether unknown alleles could be detected.

MPSeqM results for control DNA with known isolates were used to determine the best number of individual samples in one bulk. In the Mix_LM control containing DNA of 71 isolates, many alleles (>20%) present in one isolate only were not detected. This value dropped to around 6% in the bulk with 32 leaf spots and, more importantly, 100% of alleles encoding non-RIPped sequences were detected. Lack of detection of a RIPped allele present at low frequency is not a problem if the deletions (i.e. lack of amplification) can be accurately detected, because both RIPped and deleted alleles correspond to virulent versions of the gene. Altogether, comparison of pools of 32 or 96 leaf spots indicated a better detection of alleles at low frequencies with pools of 32 leaf spots. Therefore, pools of 32 leaf spots were selected to assess the MPSeqM on unknown samples.

The use of a dilution series with varying proportions of DNA of two isolates differing by the presence or absence of some avirulence genes allowed us to quantify deletions. The data (number of reads for a given avirulence gene) were first normalized with the number of reads for a single-copy, conserved gene. Here the *Actin* gene was chosen as it is reliable for species-specific identification of *L. maculans* within the *Leptosphaeria* species complex (Vincenot et al., 2009; Voigt et al., 2005). The normalized read numbers for the deleted genes in the dilution series correlated with the expected proportions. Using the dilution curves, the proportions of deleted isolates in known bulks were accurately quantified for either high (*AvrLm1*), low (*AvrLm11*), or intermediate (*AvrLm4-7*) proportions of deletion events.

Unexpected OTUs were detected at low frequencies in bulks with known *AvrLm* allele composition, corresponding to artefacts generated at the PCR or sequencing steps. The proportions of such reads in controls were used to establish the thresholds under which MiSeq reads were suppressed. Once these thresholds were applied, the datasets from field samples comprised a reduced number of allelic variants per gene, but still numerous unknown alleles. We defined rules to select reliable alleles among these new sequences. Only unknown sequences consistently present in the three replicates of one field sample and in a few samples only were kept, allowing us to identify 206 new alleles including 74 new RIPped alleles. Although not exhaustive yet, we experimentally confirmed the reality of at least some new alleles by cloning PCR products from one field sample and subsequent Sanger sequencing. Depending on the mutations present in these new alleles, the inference of the corresponding phenotype (virulent or avirulent) was more or less easy. RIPped sequences, sequences with frameshift or stop codons were considered virulent. Other variants may display silent mutations or mutations without impact on the mature protein and thus, were easily converted into avirulent or virulent alleles. However, a non-negligible amount (9.2%) of new alleles could not be converted into a phenotype. These new alleles represented low proportions of reads per sample (1.5% on average), thus without a high impact on the survey result. If an unknown allele reaches high proportions in specific sites, it will be necessary to go back to isolation campaigns at this site, with the aim of recovering and phenotyping isolates with that allelic variant, or to assess the corresponding phenotype following the transformation of a virulent isolate with a synthetic construct displaying this new allele.

The MPSeqM tool has some limitations. Because of the rather short sequence length, only part of the full CDS length is amplified for some *AvrLm* genes, possibility preventing us from detecting mutations in the 5’ or 3’ regions of the genes. The MPSeqM tool nevertheless enabled us to estimate frequencies of virulent isolates from field samples highly correlated with those determined by pathotyping. This limitation has however to be kept in mind, and surveys based on isolation, pathotyping, and full-length *AvrLm* sequencing will still have to be done regularly (for instance, every five years) to detect new virulent alleles, enrich the sequence database, and adapt the primers to amplify new alleles. Another limitation concerns *AvrLm4-7*. Some virulent isolates toward *Rlm7* do not amplify the gene with external primers but could amplify internal parts of the gene with a sequence matching avirulent alleles (Daverdin et al., 2012; Gautier et al., 2023). Such isolates represent 6.7% of the 2017 sampling (Gautier et al., 2023) and they will be miss-identified as avirulent with the MPSeqM tool. This phenomenon could explain why the MPSeqM tool detects more avirulent isolates toward *Rlm7* than pathotyping (Figure 6).

The MPSeqM field survey revealed highly contrasted levels of polymorphism depending on the *AvrLm* gene, confirming data obtained by *AvrLm* sequencing in individual isolates, either from France (Gautier et al., 2023) or worldwide (Van de Wouw et al., 2023), with highly polymorphic genes (*AvrLm3*, *AvrLm4-7* or *AvrLmS-Lep2*) or low polymorphic ones (*AvrLm2* or *AvrLm6*). Here, by analysing a total of 1337 leaf spots in one single sequencing run, we confirmed these trends but also detected a number of new, rare alleles, most of them being site specific. This work thus revealed an unexpected diversity of *AvrLm* sequences for at least some genes. Whether this polymorphism is pre-existing at a basal level, or whether it is locally generated at each generation through meiosis (Daverdin et al., 2012), is still in question. The high number of new RIPped alleles detected for *AvrLm4-7* (66 new sequences), but not for other genes suggests RIP is still an active mechanism particularly frequent for this gene, generating new alleles every year at meiosis.

## CONCLUSION

The MPSeqM tool appears suitable for a precise characterization of *L. maculans* populations as regards virulence at eight avirulence loci, without the tedious steps of *L. maculans* isolation, in vitro growth and inoculation on a plant differential set. The MPSeqM tool also gives a detailed and precise characterisation of the diversity of *AvrLm* genes in populations present in specific situations (geographical location, specific time-point, plant genotype, plant organ). We highlighted contrasting situations between sites, with some sites displaying greater diversity but few new alleles, and others being less diverse but richer in new alleles. The MPSeqM tool thus offers new perspectives for new research questions, such as: do some plant genotypes select some allelic variants along the cropping season? How plant genotypes affect allelic variants? How different are populations present in very close or distant fields? May some resistance management strategies affect the allelic diversity of avirulence genes? Finally, this tool, associated with scorings of stem canker severity, could help bringing extensive field data to determine a “critical percentage of virulent isolates” at which the resistance is in danger of being rapidly overcome, an information currently not available for the *L. maculans* / *B. napus* pathosystem.

## ACKNOWLEDGMENTS

This work was funded by the CASDAR project C2018-11 (Atipical). We thank all contributors from the CTPS field experiments that provided infected oilseed rape leaves and Julie Noah for her contribution to leaf spot recovery. The “Effectors and Pathogenesis of *L. maculans*” group benefits from the support of Saclay Plant Sciences-SPS (ANR-17-EUR-0007). We thank the GeT Genotoul platform in Toulouse, Midi-Pyrénées for the MiSeq sequencing experiments. We are grateful to the Genotoul Bioinformatics platform in Toulouse, Midi-Pyrénées for providing help in computing and data storage.

## Data availability statement

Data available on request from the authors.

## Figure legends

**Supplementary Table S1. List and characteristics of *Leptosphaeria maculans* isolates used in this study**

**Supplementary Table S2. Genotypes of reference *Leptosphaeria maculans* isolates at eight avirulence loci**. Isolates in bold are those present in the poolof 32 *Brassica napus* leaf spots used as control.

**Supplementary Table S3. List and proportions of *AvrLm* alleles present in the Mix_LM and Mix_LSpot controls and the corresponding estimated proportions with after MiSeq sequencing.**

**Supplementary Figure S1. Experimental design for field samples analyses.** 1, Leaves at reception in the lab, cleaned and air-dried; 2, one leaf spot per leaf is cut and placed in humid conditions for pycnidia production, and 3, mono-pycnidia isolation for 4, inoculation on a plant differential set to determine the phenotype. 5, standardized leaf spot cutting (one per leaf) and long-term storage in 24-well plates. 7, a view of a full plate before storage until 8, pool DNA extraction, multiplex PCR and MiSeq sequencing

**Supplementary Figure S2. Alignment of known alleles of eight *L. maculans* avirulence genes with positions of forward and reverse MiSeq primers.** A, *AvrLm1*; B, *AvrLm2*; C, *AvrLm3*; D, *AvrLm4-7*; E, *AvrLm5-9*; F*, AvrLmS-Lep2*; G*, AvrLm6*; H, *AvrLm11*. The F-primers are located before the start for *AvrLm5-9* and *AvrLm6* and the R-primer is after the stop for *AvrLm11*.

**Supplementary Figure S3. Box plot of the frequency per sampling site of avirulent alleles for *AvrLm1,* as estimated by MiSeq sequencing in 42 pools of leaf spots.**

**Supplementary Figure S4. Proportions of the three *AvrLm2* alleles detected in 42 pools of leaf spots from 9 fields**

**Supplementary Figure S5 Maximum likelihood tree of all *AvrLm3* isoforms identified here.** Isoforms underlined in green are known avirulent isoforms while those underlined in red are known virulent isoforms.

**Supplementary Figure S6. Percentages of the 41 *AvrLm3* alleles as determined following multiplex-PCR and MiSeq sequencing of 42 field samples**

**Supplementary Figure S7. Proportions of *AvrLm5-9* alleles detected in 42 pools of leaf spots from nine fields**

**Supplementary Figure S8. Proportions of *AvrLm6* alleles detected in 42 pools of leaf spots from nine fields**

**Supplementary Figure S9. Proportions of *AvrLmS-Lep2* alleles detected in 42 pools of leaf spots from nine fields**

**Supplementary Figure S10. Relationships between percentages of each *AvrLm3* OTU per site estimated in 32 leaf spots or from the “pools of pools” (64 to 96 leaf spots). Each dot corresponds to the percentage of one OTU in one site.** A, all data; B, C and D, focus on high (>60%), low (<20%) or very low (<2) percentages, respectively. Green circle highlights OTUs that are detected at low frequencies in 24 or 32 leaf spot pools but not in 64 or 96 leaf spots. The red circle highlights OTUs that are not detected in pools of 32 leaf spots but found in pools of 96, thus probably corresponding to sequencing artefacts.

**Supplementary Figure S11. Correlation between estimates of percentage of avirulent isolates on *Rlm7* by pathotyping and with the MPSeqM tool.** Each dot corresponds to one geographical location. Samplings were done at the same place and same time but on different leaves. Pathotyping was described in Balesdent et al. (2023) and was done using around 80 isolates per site, while the genotyping was done on 3 to 6 pools of 32 leaf spots per site. The Spearman correlation coefficient is indicated.

**Supplementary Figure S12. Relative *AvrLm* allele diversity per site as described with the MpSeqM tool.** A, diversity for all genes; B, diversity for *AvrLm11* only. The radar diagrams represent, for each site, the relative proportion of alleles (no. of alleles at this site over total no. of alleles detected in the survey) averaged by the relative proportion of leaf spots analysed at this site. Dark line, overall diversity; dark grey line, diversity for “new” alleles; pale grey line, diversity for site-specific new alleles. Sites as follows: Co, Coullons; Ja, Janville; Th1, Thiverval-Grignon field 1; Th2, Thiverval-Grignon field 2; Boi, Boisseau; Er, Erdre-en-Anjou; Bou, Bouges-le-Château; Fr, Frelinghein; LeR, Le Rheu

**Supplementary Text T1. Parameters used for the Mothur pipeline.**

## Notes

### Competing Interest Statement

The authors have declared no competing interest.

### Summary of Updates

(1) Mainly,an error about the B. napus cultivar used in the field was corrected (cv. Westar instead of Drakkar, as described in Balesdent et al, 2023) (2)"environmental sample" replaced by "Field sample" (3) Criteria for primer design moved from results to Materials and methods (4) new, preliminary results about the molecular validation of new alleles added (5) discussion was slightly shortened and a few grammatical errors were corrected

