## Supplementary material for "MPSeqM, a tool combining multiplex PCR and high-throughput sequencing to study the polymorphism of eight *Leptosphaeria maculans* avirulence genes and its application to field surveys in France"

**Supplementary Table S1. List and characteristics of *Leptosphaeria maculans* isolates used in this study**

| Isolate ID | Country of origin | Associated host | Associated cultivar | Region of origin | City | Isolation date | Reference |
|---|---|---|---|---|---|---|---|
| **EPH17.001**[1] | France | *Brassica napus* L. | Falcon | Seine-et-Marne (77) | Verneuil l'Etang | 2017 | Gautier et al.,2023 |
| **EPH17.004** | France | *Brassica napus* L. | Falcon | Seine-et-Marne (77) | Verneuil l'Etang | 2017 | Gautier et al.,2023 |
| **EPH17.010** | France | *Brassica napus* L. | Falcon | Seine-et-Marne (77) | Verneuil l'Etang | 2017 | Gautier et al.,2023 |
| **EPH17.016** | France | *Brassica napus* L. | Falcon | Seine-et-Marne (77) | Verneuil l'Etang | 2017 | Gautier et al.,2023 |
| **EPH17.019** | France | *Brassica napus* L. | Falcon | Seine-et-Marne (77) | Verneuil l'Etang | 2017 | Gautier et al.,2023 |
| EPH17.029 | France | *Brassica napus* L. | Falcon | Seine-et-Marne (77) | Verneuil l'Etang | 2017 | Gautier et al.,2023 |
| **EPH17.066** | France | *Brassica napus* L. | Falcon | Nord (59) | Prémesques | 2017 | Gautier et al.,2023 |
| **EPH17.071** | France | *Brassica napus* L. | Falcon | Nord (59) | Prémesques | 2017 | Gautier et al.,2023 |
| **EPH17.084** | France | *Brassica napus* L. | Falcon | Nord (59) | Prémesques | 2017 | Gautier et al.,2023 |
| **EPH17.102** | France | *Brassica napus* L. | Falcon | Loir-et-Cher (41) | Blois | 2017 | Gautier et al.,2023 |
| **EPH17.105** | France | *Brassica napus* L. | Falcon | Loir-et-Cher (41) | Blois | 2017 | Gautier et al.,2023 |
| **EPH17.108** | France | *Brassica napus* L. | Falcon | Loir-et-Cher (41) | Blois | 2017 | Gautier et al.,2023 |
| EPH17.111 | France | *Brassica napus* L. | Falcon | Loir-et-Cher (41) | Blois | 2017 | Gautier et al.,2023 |
| **EPH17.114** | France | *Brassica napus* L. | Falcon | Loir-et-Cher (41) | Blois | 2017 | Gautier et al.,2023 |
| **EPH17.117** | France | *Brassica napus* L. | Falcon | Loir-et-Cher (41) | Blois | 2017 | Gautier et al.,2023 |
| **EPH17.120** | France | *Brassica napus* L. | Falcon | Loir-et-Cher (41) | Blois | 2017 | Gautier et al.,2023 |
| EPH17.124 | France | *Brassica napus* L. | Falcon | Loir-et-Cher (41) | Blois | 2017 | Gautier et al.,2023 |
| **EPH17.127** | France | *Brassica napus* L. | Falcon | Loir-et-Cher (41) | Blois | 2017 | Gautier et al.,2023 |
| EPH17.130 | France | *Brassica napus* L. | Falcon | Loir-et-Cher (41) | Blois | 2017 | Gautier et al.,2023 |
| **EPH17.133** | France | *Brassica napus* L. | Falcon | Loir-et-Cher (41) | Blois | 2017 | Gautier et al.,2023 |
| **EPH17.142** | France | *Brassica napus* L. | Falcon | Loir-et-Cher (41) | Blois | 2017 | Gautier et al.,2023 |
| EPH17.145 | France | *Brassica napus* L. | Falcon | Haute-Garonne (31) | Seilh | 2017 | Gautier et al.,2023 |
| EPH17.151 | France | *Brassica napus* L. | Falcon | Haute-Garonne (31) | Seilh | 2017 | Gautier et al.,2023 |

| Isolate ID | Country of origin | Associated host | Associated cultivar | Region of origin | City | Isolation date | Reference |
|---|---|---|---|---|---|---|---|
| **EPH17.157** | France | *Brassica napus* L. | Falcon | Haute-Garonne (31) | Seilh | 2017 | Gautier et al.,2023 |
| **EPH17.164** | France | *Brassica napus* L. | Falcon | Haute-Garonne (31) | Seilh | 2017 | Gautier et al.,2023 |
| EPH17.167 | France | *Brassica napus* L. | Falcon | Haute-Garonne (31) | Seilh | 2017 | Gautier et al.,2023 |
| EPH17.179 | France | *Brassica napus* L. | Falcon | Haute-Garonne (31) | Seilh | 2017 | Gautier et al.,2023 |
| **EPH17.186** | France | *Brassica napus* L. | Falcon | Haute-Garonne (31) | Seilh | 2017 | Gautier et al.,2023 |
| **EPH17.194** | France | *Brassica napus* L. | Falcon | Seine-et-Marne (77) | Verneuil l'Etang | 2017 | Gautier et al.,2023 |
| EPH17.197 | France | *Brassica napus* L. | Falcon | Seine-et-Marne (77) | Verneuil l'Etang | 2017 | Gautier et al.,2023 |
| EPH17.207 | France | *Brassica napus* L. | Falcon | Seine-et-Marne (77) | Verneuil l'Etang | 2017 | Gautier et al.,2023 |
| EPH17.216 | France | *Brassica napus* L. | Falcon | Seine-et-Marne (77) | Verneuil l'Etang | 2017 | Gautier et al.,2023 |
| **EPH17.225** | France | *Brassica napus* L. | Falcon | Seine-et-Marne (77) | Verneuil l'Etang | 2017 | Gautier et al.,2023 |
| EPH17.231 | France | *Brassica napus* L. | Falcon | Seine-et-Marne (77) | Verneuil l'Etang | 2017 | Gautier et al.,2023 |
| **EPH17.234** | France | *Brassica napus* L. | Falcon | Seine-et-Marne (77) | Verneuil l'Etang | 2017 | Gautier et al.,2023 |
| **EPH17.238** | France | *Brassica napus* L. | Falcon | Nord (59) | Prémesques | 2017 | Gautier et al.,2023 |
| **EPH17.250** | France | *Brassica napus* L. | Falcon | Nord (59) | Prémesques | 2017 | Gautier et al.,2023 |
| **EPH17.256** | France | *Brassica napus* L. | Falcon | Nord (59) | Prémesques | 2017 | Gautier et al.,2023 |
| EPH17.272 | France | *Brassica napus* L. | Falcon | Nord (59) | Prémesques | 2017 | Gautier et al.,2023 |
| **EPH17.275** | France | *Brassica napus* L. | Falcon | Nord (59) | Prémesques | 2017 | Gautier et al.,2023 |
| EPH17.278 | France | *Brassica napus* L. | Falcon | Nord (59) | Prémesques | 2017 | Gautier et al.,2023 |
| **EPH17.287** | France | *Brassica napus* L. | Falcon | Loir-et-Cher (41) | Blois | 2017 | Gautier et al.,2023 |
| EPH17.296 | France | *Brassica napus* L. | Falcon | Loir-et-Cher (41) | Blois | 2017 | Gautier et al.,2023 |
| **EPH17.304** | France | *Brassica napus* L. | Falcon | Loir-et-Cher (41) | Blois | 2017 | Gautier et al.,2023 |
| EPH17.308 | France | *Brassica napus* L. | Falcon | Loir-et-Cher (41) | Blois | 2017 | Gautier et al.,2023 |
| EPH17.324 | France | *Brassica napus* L. | Falcon | Haute-Garonne (31) | Seilh | 2017 | Gautier et al.,2023 |
| EPH17.336 | France | *Brassica napus* L. | Falcon | Haute-Garonne (31) | Seilh | 2017 | Gautier et al.,2023 |
| EPH17.339 | France | *Brassica napus* L. | Falcon | Haute-Garonne (31) | Seilh | 2017 | Gautier et al.,2023 |

| Isolate ID | Country of origin | Associated host | Associated cultivar | Region of origin | City | Isolation date | Reference |
|---|---|---|---|---|---|---|---|
| JN2(v23.1.2) | Lab isolate | na | na | na | na | na | Balesdent et al, 2002 |
| **JN3 (v23.1.3)** | Lab isolate (Sequence reference) | na | na | na | na | na | Balesdent et al, 2002 Rouxel et al, 2011 |
| BBA-62908 | Germany | nd ("B. vulgaris") | na | Unknown | Unknown | 1966 | Rouxel et al, 2013 |
| CB-G3.5 | France- | *Brassica napus* L. | Drakkar | Yvelines (78) | Thiverval-Grignon | 2000 | Balesdent et al, 2006 |
| CB-i2-9 | France- | *Brassica napus* L. | Drakkar | Ille-et-Vilaine (35) | Le Rheu | 2000 | Balesdent et al, 2006 |
| IBCN14 | Australia | *Brassica napus* L. | Unknown | South Australia | Mundulla | 1988 | Balesdent et al, 2005 |
| IBCN18 | Australia | *Brassica napus* L | Unknown (Stubble) | Victoria | | 1988 | Balesdent et al, 2005 |
| PHW1245 (IBCN74) | France | *Brassica oleracea* | Unknown | Unknown | Unknown | Unknown (before 1980) | Balesdent et al, 2005 |
| IBCN75 | Australia | *Brassica napus* L | Unknown | Western Australia | Mount Barker | 1987 | Balesdent et al, 2005 |
| IBCN76 | Australia | *Brassica napus* L | Unknown | Western Australia | Mount Barker | 1987 | Balesdent et al, 2005 |
| IBCN79 | Australia | *Brassica napus* L | Unknown | Western Australia | Mount Barker | 1987 | Balesdent et al, 2005 |
| IBCN80 | Canada | *Brassica napus* L | Unknown (Stubble) | Saskatchewan | Unknown | 1985 | Balesdent et al, 2005 |
| IBCN85 | Canada | *Brassica napus* L | Unknown (Seeds) | Ontario | Unknown | 1989 | Balesdent et al, 2005 |
| MX8-12 | France | *Brassica napus* L | MX (*Rlm6*) line | Ille-et-Vilaine (35) | Le Rheu | 1994 | Fudal et al, 2009 |
| Nz-T4 | New Zealand | *Brassica rapa* | Tina | Unknown | Unknown | Unknown | Balesdent et al, 2005 |
| OMR19 | Mexico | *Brassica oleracea* | Apex | Unknown | Unknown | 2005 | Dilmaghani et al, 2013 |
| UWAP11 | Australia | *Brassica napus* L | Unknown (Stubble) | Western Australia | Unknown | 2001 | Balesdent et al, 2005 |
| WAC4028 | Australia | *Brassica napus* L | Midas | Western Australia | Unknown | 1984 | Balesdent et al, 2005 |
| WAC4057 | Australia | *Brassica napus* L | Wesreo | Western Australia | Unknown | 1984 | Balesdent et al, 2005 |
| WAC4079 | Australia | *Brassica napus* L | Wesbell | Western Australia | Unknown | 1984 | Balesdent et al, 2005 |
| WAC7750 | Australia | Unknown | Unknown | Western Australia | Unknown | 1972 | Balesdent et al, 2005 |

| Isolate ID | Country of origin | Associated host | Associated cultivar | Region of origin | City | Isolation date | Reference |
|---|---|---|---|---|---|---|---|
| WAC7803 | Australia | *Raphanus raphanistrum* | Unknown | Western Australia | Unknown | 1973 | Balesdent et al, 2005 |
| HB10-19 | France | *Brassica nigra* | Junius | Ille et Vilaine (35) | Le Rheu | 1995 | This study |

[1] Isolates in bold are those inoculated to produce the control DNA of mixes of leaf lesions of known composition in Avr alleles

[2] na, not applicable

**SupplementaryTable S2. Genotypes of reference *Leptosphaeria maculans* isolates at eight avirulence loci. Isolates in bold are those used in the mix of 32 *Brassica napus* leaf spots used as control.**

| Isolate | *AvrLm1* | *AvrLm2* | *AvrLm3* | *AvrLm4-7* | *AvrLm5-9* | *AvrLm6* | *AvrLm11* | *AvrLmS-Lep2* |
|---|---|---|---|---|---|---|---|---|
| **EPH17.001** | *AvrLm1_del*[1] | *AvrLm2_0_A* | *AvrLm3_A_01* | *AvrLm4-7_del* | *AvrLm5-9_0_A* | *AvrLm6_0_A* | *AvrLm11_0_A* | *AvrLmS_2_B* |
| **EPH17.004** | *AvrLm1_del* | *AvrLm2_0_A* | *AvrLm3_A_01* | *AvrLm4-7_del* | *AvrLm5-9_0_A* | *AvrLm6_0_A* | *AvrLm11_0_A* | *AvrLmS_7_A* |
| **EPH17.010** | *AvrLm1_del* | *AvrLm2_0_A* | *AvrLm3_C_01* | *AvrLm4-7_rip_1* | *AvrLm5-9_0_B* | *AvrLm6_0_A* | *AvrLm11_0_A* | *AvrLmS_2_B* |
| **EPH17.016** | *AvrLm1_del* | *AvrLm2_0_A* | *AvrLm3_A_09* | *AvrLm4-7_9_A* | *AvrLm5-9_0_A* | *AvrLm6_0_A* | *AvrLm11_0_A* | *AvrLmS_2_B* |
| **EPH17.019** | *AvrLm1_del* | *AvrLm2_0_A* | *AvrLm3_A_09* | *AvrLm4-7_partial_del* | *AvrLm5-9_0_B* | *AvrLm6_0_A* | *AvrLm11_0_A* | *AvrLmS_2_F* |
| EPH17.029 | *AvrLm1_del* | *AvrLm2_0_A* | *AvrLm3_A_01* | *AvrLm4-7_del* | *AvrLm5-9_0_A* | *AvrLm6_0_A* | *AvrLm11_del* | *AvrLmS_2_B* |
| **EPH17.066** | *AvrLm1_del* | *AvrLm2_0_A* | *AvrLm3_H_02* | *AvrLm4-7_partial_del* | *AvrLm5-9_0_A* | *AvrLm6_0_A* | *AvrLm11_0_A* | *AvrLmS_4_A* |
| **EPH17.071** | *AvrLm1_rip3* | *AvrLm2_0_A* | *AvrLm3_C_01* | *AvrLm4-7_1_A* | *AvrLm5-9_0_A* | *AvrLm6_0_A* | *AvrLm11_0_A* | *AvrLmS_6_A* |
| **EPH17.084** | *AvrLm1_del* | *AvrLm2_0_A* | *AvrLm3_A_01* | *AvrLm4-7_5_A* | *AvrLm5-9_0_B* | *AvrLm6_0_A* | *AvrLm11_0_A* | *AvrLmS_2_B* |
| **EPH17.102** | *AvrLm1_del* | *AvrLm2_1_A* | *AvrLm3_H_02* | *AvrLm4-7_del* | *AvrLm5-9_0_A* | *AvrLm6_0_A* | *AvrLm11_0_A* | *AvrLmS_2_B* |
| **EPH17.105** | *AvrLm1_del* | *AvrLm2_0_A* | *AvrLm3_A_09* | *AvrLm4-7_6_A* | *AvrLm5-9_0_A* | *AvrLm6_0_A* | *AvrLm11_0_A* | *AvrLmS_2_A* |
| **EPH17.108** | *AvrLm1_del* | *AvrLm2_0_A* | *AvrLm3_A_09* | *AvrLm4-7_rip_2* | *AvrLm5-9_0_B* | *AvrLm6_0_A* | *AvrLm11_0_A* | *AvrLmS_2_B* |
| EPH17.111 | *AvrLm1_del* | *AvrLm2_0_A* | *AvrLm3_A_09* | *AvrLm4-7_5_A* | *AvrLm5-9_0_A* | *AvrLm6_0_A* | *AvrLm11_0_A* | *AvrLmS_0_A* |
| **EPH17.114** | *AvrLm1_del* | *AvrLm2_0_A* | *AvrLm3_A_09* | *AvrLm4-7_7_A* | *AvrLm5-9_0_A* | *AvrLm6_0_A* | *AvrLm11_0_A* | *AvrLmS_2_B* |
| **EPH17.117** | *AvrLm1_del* | *AvrLm2_0_A* | *AvrLm3_A_09* | *AvrLm4-7_2_A* | *AvrLm5-9_0_A* | *AvrLm6_0_A* | *AvrLm11_0_A* | *AvrLmS_2_B* |
| **EPH17.120** | *AvrLm1_del* | *AvrLm2_0_A* | *AvrLm3_A_09* | *AvrLm4-7_del_AA_A* | *AvrLm5-9_0_A* | *AvrLm6_0_A* | *AvrLm11_del* | *AvrLmS_2_B* |
| EPH17.124 | *AvrLm1_del* | *AvrLm2_0_A* | *AvrLm3_H_02* | *AvrLm4-7_partial_del* | *AvrLm5-9_0_A* | *AvrLm6_0_A* | *AvrLm11_0_A* | *AvrLmS_2_B* |
| EPH17.127 | *AvrLm1_del* | *AvrLm2_0_A* | *AvrLm3_A_01* | *AvrLm4-7_1_A* | *AvrLm5-9_0_A* | *AvrLm6_0_A* | *AvrLm11_0_A* | *AvrLmS_2_B* |
| EPH17.130 | *AvrLm1_del* | *AvrLm2_0_A* | *AvrLm3_A_01* | *AvrLm4-7_ripdup_A* | *AvrLm5-9_0_A* | *AvrLm6_0_A* | *AvrLm11_0_A* | *AvrLmS_2_B* |
| **EPH17.133** | *AvrLm1_del* | *AvrLm2_0_A* | *AvrLm3_H_02* | *AvrLm4-7_del* | *AvrLm5-9_0_C* | *AvrLm6_0_A* | *AvrLm11_0_A* | *AvrLmS_2_D* |
| **EPH17.142** | *AvrLm1_del* | *AvrLm2_0_A* | *AvrLm3_A_01* | *AvrLm4-7_8_A* | *AvrLm5-9_0_A* | *AvrLm6_0_A* | *AvrLm11_0_A* | *AvrLmS_1_A* |
| EPH17.145 | *AvrLm1_del* | *AvrLm2_0_A* | *AvrLm3_A_01* | *AvrLm4-7_2_A* | *AvrLm5-9_0_A* | *AvrLm6_0_A* | *AvrLm11_0_A* | *AvrLmS_7_A* |
| EPH17.151 | *AvrLm1_del* | *AvrLm2_0_A* | *AvrLm3_A_09* | *AvrLm4-7_1_A* | *AvrLm5-9_0_A* | *AvrLm6_0_A* | *AvrLm11_0_A* | *AvrLmS_0_A* |
| **EPH17.157** | *AvrLm1_del* | *AvrLm2_1_A* | *AvrLm3_A_09* | *AvrLm4-7_2_A* | *AvrLm5-9_0_C* | *AvrLm6_0_A* | *AvrLm11_0_A* | *AvrLmS_2_B* |
| EPH17.164 | *AvrLm1_del* | *AvrLm2_0_A* | *AvrLm3_C_01* | *AvrLm4-7_2_A* | *AvrLm5-9_0_B* | *AvrLm6_0_A* | *AvrLm11_0_A* | *AvrLmS_2_B* |
| EPH17.167 | *AvrLm1_del* | *AvrLm2_0_A* | *AvrLm3_A_01* | *AvrLm4-7_1_A* | *AvrLm5-9_0_A* | *AvrLm6_0_A* | *AvrLm11_0_A* | *AvrLmS_4_A* |
| EPH17.179 | *AvrLm1_del* | *AvrLm2_0_A* | *AvrLm3_A_01* | *AvrLm4-7_1_A* | *AvrLm5-9_0_A* | *AvrLm6_0_A* | *AvrLm11_0_A* | *AvrLmS_1_A* |

| | | | | | | | | |
|---|---|---|---|---|---|---|---|---|
| **EPH17.186** | *AvrLm1_del* | *AvrLm2_0_A* | *AvrLm3_A_09* | *AvrLm4-7_1_A* | *AvrLm5-9_0_A* | *AvrLm6_0_A* | *AvrLm11_0_A* | *AvrLmS_2_C* |
| **EPH17.194** | *AvrLm1_del* | *AvrLm2_0_A* | *AvrLm3_A_01* | *AvrLm4-7_10_A* | *AvrLm5-9_0_A* | *AvrLm6_0_A* | *AvrLm11_0_A* | *AvrLmS_2_B* |
| EPH17.197 | *AvrLm1_del* | *AvrLm2_0_A* | *AvrLm3_A_09* | *AvrLm4-7_del* | *AvrLm5-9_0_A* | *AvrLm6_0_A* | *AvrLm11_0_A* | *AvrLmS_2_B* |
| EPH17.207 | *AvrLm1_del* | *AvrLm2_0_A* | *AvrLm3_A_01* | *AvrLm4-7_2_A* | *AvrLm5-9_0_A* | *AvrLm6_0_A* | *AvrLm11_0_A* | *AvrLmS_0_A* |
| EPH17.216 | *AvrLm1_del* | *AvrLm2_0_A* | *AvrLm3_C_01* | *AvrLm4-7_del* | *AvrLm5-9_0_B* | *AvrLm6_0_A* | *AvrLm11_0_A* | *AvrLmS_2_B* |
| **EPH17.225** | *AvrLm1_del* | *AvrLm2_0_A* | *AvrLm3_I_01* | *AvrLm4-7_partial_del* | *AvrLm5-9_0_A* | *AvrLm6_0_A* | *AvrLm11_0_A* | *AvrLmS_2_C* |
| EPH17.231 | *AvrLm1_del* | *AvrLm2_0_A* | *AvrLm3_A_09* | *AvrLm4-7_del* | *AvrLm5-9_0_A* | *AvrLm6_0_A* | *AvrLm11_0_A* | *AvrLmS_4_A* |
| **EPH17.234** | *AvrLm1_del* | *AvrLm2_0_A* | *AvrLm3_J_03* | *AvrLm4-7_del* | *AvrLm5-9_0_C* | *AvrLm6_0_A* | *AvrLm11_0_A* | *AvrLmS_2_B* |
| **EPH17.238** | *AvrLm1_0_A* | *AvrLm2_1_A* | *AvrLm3_A_09* | *AvrLm4-7_1_A* | *AvrLm5-9_0_A* | *AvrLm6_0_A* | *AvrLm11_0_A* | *AvrLmS_3_A* |
| **EPH17.250** | *AvrLm1_del* | *AvrLm2_0_A* | *AvrLm3_A_07* | *AvrLm4-7_2_A* | *AvrLm5-9_0_A* | *AvrLm6_0_A* | *AvrLm11_del* | *AvrLmS_2_B* |
| **EPH17.256** | *AvrLm1_0_A* | *AvrLm2_0_A* | *AvrLm3_C_01* | *AvrLm4-7_rip_3* | *AvrLm5-9_0_A* | *AvrLm6_0_A* | *AvrLm11_0_A* | *AvrLmS_5_A* |
| EPH17.272 | *AvrLm1_del* | *AvrLm2_0_A* | *AvrLm3_A_01* | *AvrLm4-7_5_A* | *AvrLm5-9_0_C* | *AvrLm6_0_A* | *AvrLm11_0_A* | *AvrLmS_2_D* |
| **EPH17.275** | *AvrLm1_del* | *AvrLm2_1_A* | *AvrLm3_C_01* | *AvrLm4-7_1_A* | *AvrLm5-9_0_A* | *AvrLm6_3_A* | *AvrLm11_0_A* | *AvrLmS_2_B* |
| EPH17.278 | *AvrLm1_0_A* | *AvrLm2_0_A* | *AvrLm3_A_09* | *AvrLm4-7_1_A* | *AvrLm5-9_0_A* | *AvrLm6_0_A* | *AvrLm11_0_A* | *AvrLmS_2_B* |
| **EPH17.287** | *AvrLm1_del* | *AvrLm2_0_A* | *AvrLm3_A_09* | *AvrLm4-7_del* | *AvrLm5-9_0_A* | *AvrLm6_0_A* | *AvrLm11_0_A* | *AvrLmS_2_G* |
| EPH17.296 | *AvrLm1_del* | *AvrLm2_0_A* | *AvrLm3_A_01* | *AvrLm4-7_del* | *AvrLm5-9_0_C* | *AvrLm6_0_A* | *AvrLm11_0_A* | *AvrLmS_1_A* |
| **EPH17.304** | *AvrLm1_del* | *AvrLm2_0_A* | *AvrLm3_J_04* | *AvrLm4-7_del* | *AvrLm5-9_0_A* | *AvrLm6_0_A* | *AvrLm11_0_A* | *AvrLmS_0_A* |
| EPH17.308 | *AvrLm1_del* | *AvrLm2_0_A* | *AvrLm3_A_09* | *AvrLm4-7_1_A* | *AvrLm5-9_0_B* | *AvrLm6_0_A* | *AvrLm11_del* | *AvrLmS_7_A* |
| EPH17.324 | *AvrLm1_del* | *AvrLm2_0_A* | *AvrLm3_A_05* | *AvrLm4-7_2_A* | *AvrLm5-9_0_A* | *AvrLm6_0_A* | *AvrLm11_0_A* | *AvrLmS_1_A* |
| EPH17.336 | *AvrLm1_0_A* | *AvrLm2_0_A* | *AvrLm3_A_05* | *AvrLm4-7_del* | *AvrLm5-9_0_B* | *AvrLm6_0_A* | *AvrLm11_0_A* | *AvrLmS_0_A* |
| EPH17.339 | *AvrLm1_del* | *AvrLm2_0_A* | *AvrLm3_A_16* | *AvrLm4-7_2_A* | *AvrLm5-9_0_A* | *AvrLm6_0_A* | *AvrLm11_0_A* | *AvrLmS_2_B* |
| JN2(v23.1.2) | *AvrLm1_del* | *AvrLm2_0_A* | *AvrLm3_A_09* | *AvrLm4-7_1_A* | *AvrLm5-9_0_A* | *AvrLm6_0_A* | *AvrLm11_0_A* | *AvrLmS_1_A* |
| **JN3 (v23.1.3)** | *AvrLm1_0_A* | *AvrLm2_0_A* | *AvrLm3_A_01* | *AvrLm4-7_0_A* | *AvrLm5-9_0_A* | *AvrLm6_0_A* | *AvrLm11_0_A* | *AvrLmS_0_A* |
| BBA-62908 | *AvrLm1_0_A* | *AvrLm2_3_A* | *AvrLm3_G_1* | *AvrLm4-7_4_A* | *AvrLm5-9_1_A* | *AvrLm6_1_A* | *AvrLm11_1_A* | *AvrLmS_8_B* |
| CB-G3.5 | *AvrLm1-rip2* | *AvrLm2_0_A* | *AvrLm3_A_01* | *AvrLm4-7_1_A* | *AvrLm5-9_0_A* | *AvrLm6_0_A* | *AvrLm11_0_A* | *AvrLmS_0_A* |
| CB-i2-9 | *AvrLm1-rip1* | *AvrLm2_0_A* | *AvrLm3_A_09* | *AvrLm4-7_1_A* | *AvrLm5-9_0_A* | *AvrLm6_0_A* | *AvrLm11_0_A* | *AvrLmS_2_B* |
| IBCN14 | *AvrLm1_del* | *AvrLm2_0_A* | *AvrLm3_J_02* | *AvrLm4-7_del* | *AvrLm5-9_0_A* | *AvrLm6_1_A* | *AvrLm11_del* | *AvrLmS_2_B* |
| IBCN18 | *AvrLm1_1_A* | *AvrLm2_3_A* | *AvrLm3_P_01* | *AvrLm4-7_0_A* | *AvrLm5-9_4_A* | *AvrLm6_del* | *AvrLm11-0+2-dup* | *AvrLmS_7_A* |
| PHW1245 (IBCN74) | *AvrLm1_1_B* | *AvrLm2_4_A* | *AvrLm3_F_02* | *AvrLm4-7_3_A* | *AvrLm5-9_1_A* | *AvrLm6_2_A* | *AvrLm11_2_A* | *AvrLmS_8_B* |

| | | | | | | | | |
|---|---|---|---|---|---|---|---|---|
| IBCN75 | *AvrLm1_0_A* | *AvrLm2_0_A* | *AvrLm3_J_03* | *AvrLm4-7_del* | *AvrLm5-9_1_A* | *AvrLm6_1_A* | *AvrLm11_0_A* | *AvrLmS_7_A* |
| IBCN76 | *AvrLm1_0_A* | *AvrLm2_1_A* | *AvrLm3_A_01* | *AvrLm4-7_del* | *AvrLm5-9_0_A* | *AvrLm6_0_A* | *AvrLm11_0_A* | *AvrLmS_2_B* |
| IBCN79 | *AvrLm1_0_A* | *AvrLm2_1_A* | *AvrLm3_A_09* | *AvrLm4-7_del* | *AvrLm5-9_0_A* | *AvrLm6_0_A* | *AvrLm11_0_A* | *AvrLmS_7_A* |
| IBCN80 | *AvrLm1_del* | *AvrLm2_3_A* | *AvrLm3_D_01* | *AvrLm4-7_del* | *AvrLm5-9_1_A* | *AvrLm6_1_A* | *AvrLm11_1_A* | *AvrLmS_8_A* |
| IBCN85 | *AvrLm1_del* | *AvrLm2_2_A* | *AvrLm3_A_09* | *AvrLm4-7_2_A* | *AvrLm5-9_0_A* | *AvrLm6_1_A* | *AvrLm11_0_A* | *AvrLmS_8_B* |
| MX8-12 | *AvrLm1_0_A* | *AvrLm2_1_A* | *AvrLm3_A_01* | *AvrLm4-7_2_A* | *AvrLm5-9_0_B* | *AvrLm6_rip_1* | *AvrLm11_del* | *AvrLmS_4_B* |
| Nz-T4 | *AvrLm1_del* | *AvrLm2_0_A* | *AvrLm3_I_01* | *AvrLm4-7_rip* | *AvrLm5-9_0_A* | *AvrLm6_1_A* | *AvrLm11_0_A* | *AvrLmS_2_A* |
| OMR19 | *AvrLm1_0_A* | *AvrLm2_3_A* | *AvrLm3_F_02* | *AvrLm4-7_3_A* | *AvrLm5-9_1_A* | *AvrLm6_1_A* | *AvrLm11_1_A* | *AvrLmS_8_B* |
| UWAP11 | *AvrLm1_0_A* | *AvrLm2_1_A* | *AvrLm3_A_17* | *AvrLm4-7_1_A* | *AvrLm5-9_5_A* | *AvrLm6_0_A* | *AvrLm11_0_A* | *AvrLmS_2_B* |
| WAC4028 | *AvrLm1_del* | *AvrLm2_1_A* | *AvrLm3_A_09* | *AvrLm4-7_del* | *AvrLm5-9_0_A* | *AvrLm6_3_A* | *AvrLm11_0_A* | *AvrLmS_2_E* |
| WAC4057 | *AvrLm1_0_A* | *AvrLm2_0_A* | *AvrLm3_A_09* | *AvrLm4-7_del* | *AvrLm5-9_0_A* | *AvrLm6_1_A* | *AvrLm11_0_A* | *AvrLmS_2_B* |
| WAC4079 | *AvrLm1_0_A* | *AvrLm2_2_A* | *AvrLm3_K_01* | *AvrLm4-7_del* | *AvrLm5-9_0_A* | *AvrLm6_1_A* | *AvrLm11_0_A* | *AvrLmS_2_B* |
| WAC7750 | *AvrLm1_0_A* | *AvrLm2_2_A* | *AvrLm3_A_01* | *AvrLm4-7_0_A* | *AvrLm5-9_1_A* | *AvrLm6_1_A* | *AvrLm11_0_A* | *AvrLmS_2_B* |
| WAC7803 | *AvrLm1_del* | *AvrLm2_0_A* | *AvrLm3_K_01* | *AvrLm4-7_del* | *AvrLm5-9_3_A* | *AvrLm6_1_A* | *AvrLm11_0_A* | *AvrLmS_2_A* |
| HB10-19 | *AvrLm1_del* | *AvrLm2_X1* | *AvrLm3_del* | *AvrLm4-7_del* | *AvrLm5-9_del* | *AvrLm6_X1* | *AvrLm11_X1* | *AvrLmS_X1_A* |

[1] allele names as defined in Gautier et al., 2023. *del*, deleted allele.

**Supplementary Table S3.** List and proportions of *AvrLm* alleles present in the Mix_Lm (left) and Mix_LSpot (right) controls and the corresponding estimated proportions after MiSeq sequencing.

| Gene | Allele | Expected proportion in the Mix_Lm control | Observed proportion |
|---|---|---|---|
| *AvrLm1* | *AvrLm1_0_A*[1] | 0.2113 | 0.2892 |
| *AvrLm1* | *AvrLm1_1_A* | 0.0141 | 0.0012 |
| *AvrLm1* | *AvrLm1_2_B* | 0.0141 | 0.0018 |
| *AvrLm1* | *AvrLm1_rip_1* | 0.0141 | 0.0000 |
| *AvrLm1* | *AvrLm1_rip_2* | 0.0141 | 0.0000 |
| *AvrLm1* | *AvrLm1_rip_3* | 0.0141 | 0.0000 |
| *AvrLm1* | *AvrLm1_del* | 0.7183 | 0.7079 |
| *AvrLm2* | *AvrLm2_0_A* | 0.7465 | 0.8192 |
| *AvrLm2* | *AvrLm2_1_A* | 0.1268 | 0.1031 |
| *AvrLm2* | *AvrLm2_2_A* | 0.0423 | 0.0307 |
| *AvrLm2* | *AvrLm2_3_A* | 0.0563 | 0.0400 |
| *AvrLm2* | *AvrLm2_4_A* | 0.0141 | 0.0013 |
| *AvrLm2* | *AvrLm2_X1_A* | 0.0141 | 0.0057 |
| *AvrLm3* | *AvrLm3_A_01* | 0.5915 | 0.6552 |
| *AvrLm3* | *AvrLm3_A_05* | 0.0282 | 0.0381 |
| *AvrLm3* | *AvrLm3_A_07* | 0.0141 | 0.0160 |
| *AvrLm3* | *AvrLm3_A_16* | 0.0141 | 0.0129 |
| *AvrLm3* | *AvrLm3_A_17* | 0.0141 | 0.0068 |
| *AvrLm3* | *AvrLm3_C_01* | 0.0845 | 0.1340 |
| *AvrLm3* | *AvrLm3_D_01* | 0.0141 | 0.0051 |
| *AvrLm3* | *AvrLm3_F_02* | 0.0282 | 0.0113 |
| *AvrLm3* | *AvrLm3_G_01* | 0.0141 | 0.0055 |

| Gene | Allele | Expected proportion in the Mix_LSpot control | Observed proportion |
|---|---|---|---|
| *AvrLm1* | *AvrLm1-0_A* | 0.0938 | 0.1545 |
| *AvrLm1* | *AvrLm1_del* | 0.8750 | 0.8455 |
| *AvrLm1* | *AvrLm1_rip3* | 0.0313 | 0.0000 |
| *AvrLm2* | *AvrLm2_0_A* | 0.8750 | 0.9171 |
| *AvrLm2* | *AvrLm2_1_A* | 0.1250 | 0.0829 |
| *AvrLm3* | *AvrLm3_A_1* | 0.5938 | 0.7070 |
| *AvrLm3* | *AvrLm3_A_5* | 0.0313 | 0.0141 |
| *AvrLm3* | *AvrLm3_A_7* | 0.0313 | 0.0121 |
| *AvrLm3* | *AvrLm3_C_01* | 0.1563 | 0.1914 |
| *AvrLm3* | *AvrLm3_H_2* | 0.0938 | 0.0409 |
| *AvrLm3* | *AvrLm3_I_1* | 0.0313 | 0.0040 |
| *AvrLm3* | *AvrLm3_J_4* | 0.0625 | 0.0133 |
| *AvrLm4-7* | *AvrLm4-7_0_A* | 0.0313 | 0.0285 |
| *AvrLm4-7* | *AvrLm4-7_1_A* | 0.1563 | 0.1991 |
| *AvrLm4-7* | *AvrLm4-7_10_A* | 0.0313 | 0.0233 |
| *AvrLm4-7* | *AvrLm4-7_2_A* | 0.2031 | 0.1736 |
| *AvrLm4-7* | *AvrLm4-7_5_A* | 0.0313 | 0.0241 |
| *AvrLm4-7* | *AvrLm4-7_6_A* | 0.0313 | 0.0371 |
| *AvrLm4-7* | *AvrLm4-7_7_A* | 0.0313 | 0.0072 |
| *AvrLm4-7* | *AvrLm4-7_8_A* | 0.0313 | 0.0354 |
| *AvrLm4-7* | *AvrLm4-7_9_A* | 0.0625 | 0.0176 |
| *AvrLm4-7* | *AvrLm4-7_del* | 0.2500 | 0.3031 |

| | | | |
|---|---|---|---|
| *AvrLm3* | *AvrLm3_H_02* | 0.0563 | 0.0281 |
| *AvrLm3* | *AvrLm3_I_01* | 0.0282 | 0.0150 |
| *AvrLm3* | *AvrLm3_J_02* | 0.0141 | 0.0084 |
| *AvrLm3* | *AvrLm3_J_04* | 0.0704 | 0.0334 |
| *AvrLm3* | *AvrLm3_P_01* | 0.0141 | 0.0006 |
| *AvrLm3* | *AvrLm3_del* | 0.0141 | 0.0000 |
| *AvrLm4-7* | *AvrLm4-7_0_A* | 0.0493 | 0.0139 |
| *AvrLm4-7* | *AvrLm4-7_1_A* | 0.1972 | 0.1911 |
| *AvrLm4-7* | *AvrLm4-7_2_A* | 0.1901 | 0.1617 |
| *AvrLm4-7* | *AvrLm4-7_3_A* | 0.0282 | 0.0091 |
| *AvrLm4-7* | *AvrLm4-7_4_A* | 0.0141 | 0.0075 |
| *AvrLm4-7* | *AvrLm4-7_5_A* | 0.0423 | 0.0299 |
| *AvrLm4-7* | *AvrLm4-7_6_A* | 0.0141 | 0.0152 |
| *AvrLm4-7* | *AvrLm4-7_7_A* | 0.0141 | 0.0094 |
| *AvrLm4-7* | *AvrLm4-7_8_A* | 0.0141 | 0.0126 |
| *AvrLm4-7* | *AvrLm4-7_9_A* | 0.0282 | 0.0114 |
| *AvrLm4-7* | *AvrLm4-7_10_A* | 0.0141 | 0.0130 |
| *AvrLm4-7* | *AvrLm4-7_del* | 0.2113 | 0.3964 |
| *AvrLm4-7* | *AvrLm4-7_del_AA_1_A* | 0.0141 | 0.0134 |
| *AvrLm4-7* | *AvrLm4-7_rip_1* | 0.0141 | 0.0000 |
| *AvrLm4-7* | *AvrLm4-7_rip_100* | 0.0141 | 0.0072 |
| *AvrLm4-7* | *AvrLm4-7_rip_101* | 0.0070 | 0.0000 |
| *AvrLm4-7* | *AvrLm4-7_rip_109* | 0.0141 | 0.0036 |
| *AvrLm4-7* | *AvrLm4-7_rip_110* | 0.0141 | 0.0004 |
| *AvrLm4-7* | *AvrLm4-7_rip_96* | 0.0141 | 0.0000 |
| *AvrLm4-7* | *AvrLm4-7_rip_97* | 0.0070 | 0.0000 |
| *AvrLm4-7* | *AvrLm4-7_ripdup_A* | 0.0141 | 0.0000 |
| *AvrLm4-7* | *AvrLm4-7_ripdup_B* | 0.0141 | 0.0000 |
| *AvrLm4-7* | *AvrLm4-7_ripdup_D* | 0.0282 | 0.0000 |
| *AvrLm4-7* | *AvrLm4-7_ripdup_H* | 0.0141 | 0.0000 |

| | | | |
|---|---|---|---|
| *AvrLm4-7* | *AvrLm4-7_del_AA_A* | 0.0313 | 0.0282 |
| *AvrLm4-7* | *AvrLm4-7_rip_1* | 0.0313 | 0.0000 |
| *AvrLm4-7* | *AvrLm4-7_rip_101* | 0.0156 | 0.0000 |
| *AvrLm4-7* | *AvrLm4-7_rip_109* | 0.0313 | 0.0195 |
| *AvrLm4-7* | *AvrLm4-7_rip_110* | 0.0313 | 0.0021 |
| *AvrLm5-9* | *AvrLm5-9_0_A* | 0.8125 | 0.7572 |
| *AvrLm5-9* | *AvrLm5-9_0_B* | 0.1875 | 0.2403 |
| *AvrLm6* | *AvrLm6_0* | 0.9688 | 0.9915 |
| *AvrLm6* | *AvrLm6_1* | 0.0313 | 0.0083 |
| *AvrLm11* | *AvrLm11_0_A* | 0.9375 | 0.9766 |
| *AvrLm11* | *AvrLm11_del* | 0.0625 | 0.0096 |
| *AvrLmS* | *AvrLmS_0_A* | 0.0625 | 0.0496 |
| *AvrLmS* | *AvrLmS_1_A* | 0.0625 | 0.0496 |
| *AvrLmS* | *AvrLmS_2_A* | 0.0313 | 0.0401 |
| *AvrLmS* | *AvrLmS_2_B* | 0.5000 | 0.4847 |
| *AvrLmS* | *AvrLmS_2_C* | 0.0625 | 0.0543 |
| *AvrLmS* | *AvrLmS_2_D* | 0.0313 | 0.0387 |
| *AvrLmS* | *AvrLmS_2_E* | 0.0313 | 0.0326 |
| *AvrLmS* | *AvrLmS_2_F* | 0.0313 | 0.0136 |
| *AvrLmS* | *AvrLmS_2_G* | 0.0313 | 0.0387 |
| *AvrLmS* | *AvrLmS_3_A* | 0.0313 | 0.0278 |
| *AvrLmS* | *AvrLmS_4_A* | 0.0313 | 0.0441 |
| *AvrLmS* | *AvrLmS_5_A* | 0.0313 | 0.0272 |
| *AvrLmS* | *AvrLmS_6_A* | 0.0313 | 0.0319 |
| *AvrLmS* | *AvrLmS_7_A* | 0.0313 | 0.0109 |

| | | | |
|---|---|---|---|
| *AvrLm4-7* | *AvrLm4-7_ripdup_I* | 0.0141 | 0.0000 |
| *AvrLm5-9* | *AvrLm5-9_0_A* | 0.7320 | 0.7793 |
| *AvrLm5-9* | *AvrLm5-9_0_B* | 0.1410 | 0.1439 |
| *AvrLm5-9* | *AvrLm5-9_1_A* | 0.0704 | 0.0628 |
| *AvrLm5-9* | *AvrLm5-9_3_A* | 0.0141 | 0.0083 |
| *AvrLm5-9* | *AvrLm5-9_4_A* | 0.0141 | 0.0000 |
| *AvrLm5-9* | *AvrLm5-9_5_A* | 0.0141 | 0.0027 |
| *AvrLm5-9* | *AvrLm5-9_del* | 0.0141 | 0.0000 |
| *AvrLm6* | *AvrLm6_0_A* | 0.7606 | 0.8356 |
| *AvrLm6* | *AvrLm6_1_A* | 0.1831 | 0.1623 |
| *AvrLm6* | *AvrLm6_2_A* | 0.0141 | 0.0018 |
| *AvrLm6* | *AvrLm6_rip_1* | 0.0141 | 0.0000 |
| *AvrLm6* | *AvrLm6_X1_A* | 0.0141 | 0.0000 |
| *AvrLm6* | *AvrLm6_del* | 0.0141 | 0.0000 |
| *AvrLm11* | *AvrLm11_0_A* | 0.8310 | 0.8210 |
| *AvrLm11* | *AvrLm11_1_A* | 0.0423 | 0.0528 |
| *AvrLm11* | *AvrLm11_2_A* | 0.0141 | 0.0054 |
| *AvrLm11* | *AvrLm11_0+2_A_dup* | 0.0141 | 0.0000 |
| *AvrLm11* | *AvrLm11_del* | 0.0845 | 0.1023 |
| *AvrLm11* | *AvrLm11_X1_A* | 0.0141 | 0.0058 |
| *AvrLmS* | *AvrLmS_0_A* | 0.0986 | 0.0572 |
| *AvrLmS* | *AvrLmS_1_A* | 0.0704 | 0.0664 |
| *AvrLmS* | *AvrLmS_2_A* | 0.0423 | 0.0459 |
| *AvrLmS* | *AvrLmS_2_B* | 0.4225 | 0.5184 |
| *AvrLmS* | *AvrLmS_2_C* | 0.0282 | 0.0318 |
| *AvrLmS* | *AvrLmS_2_D* | 0.0282 | 0.0346 |
| *AvrLmS* | *AvrLmS_2_E* | 0.0282 | 0.0099 |
| *AvrLmS* | *AvrLmS_2_F* | 0.0141 | 0.0078 |
| *AvrLmS* | *AvrLmS_2_G* | 0.0141 | 0.0374 |
| *AvrLmS* | *AvrLmS_3_A* | 0.0141 | 0.0155 |
| *AvrLmS* | *AvrLmS_4_A* | 0.0423 | 0.0410 |

| | | | |
|---|---|---|---|
| *AvrLmS* | *AvrLmS_4_B* | 0.0141 | 0.0064 |
| *AvrLmS* | *AvrLmS_5_A* | 0.0141 | 0.0184 |
| *AvrLmS* | *AvrLmS_6_A* | 0.0141 | 0.0120 |
| *AvrLmS* | *AvrLmS_7_A* | 0.0845 | 0.0198 |
| *AvrLmS* | *AvrLmS_8_A* | 0.0141 | 0.0014 |
| *AvrLmS* | *AvrLmS_8_B* | 0.0423 | 0.0374 |
| *AvrLmS* | *AvrLmS_X1_A* | 0.0141 | 0.0000 |

[1] allele names as defined in Gautier et al., 2023. *del*, deleted allele.

**Supplementary Figure S1**. **Experimental design for environmental samples analyses.** 1, Leaves at reception in the lab, cleaned and air-dried; 2, one leaf spot per leaf is cut and placed in humid conditions for pycnidia production, and 3, mono-pycnidia isolation for 4, inoculation on a plant differential set to determine the phenotype. 5, standardized leaf spot cutting (one per leaf) and long term storage in 24-well plates. 7, a view of a full plate before storage until 8, pool DNA extraction, multiplex PCR and MiSeq sequencing

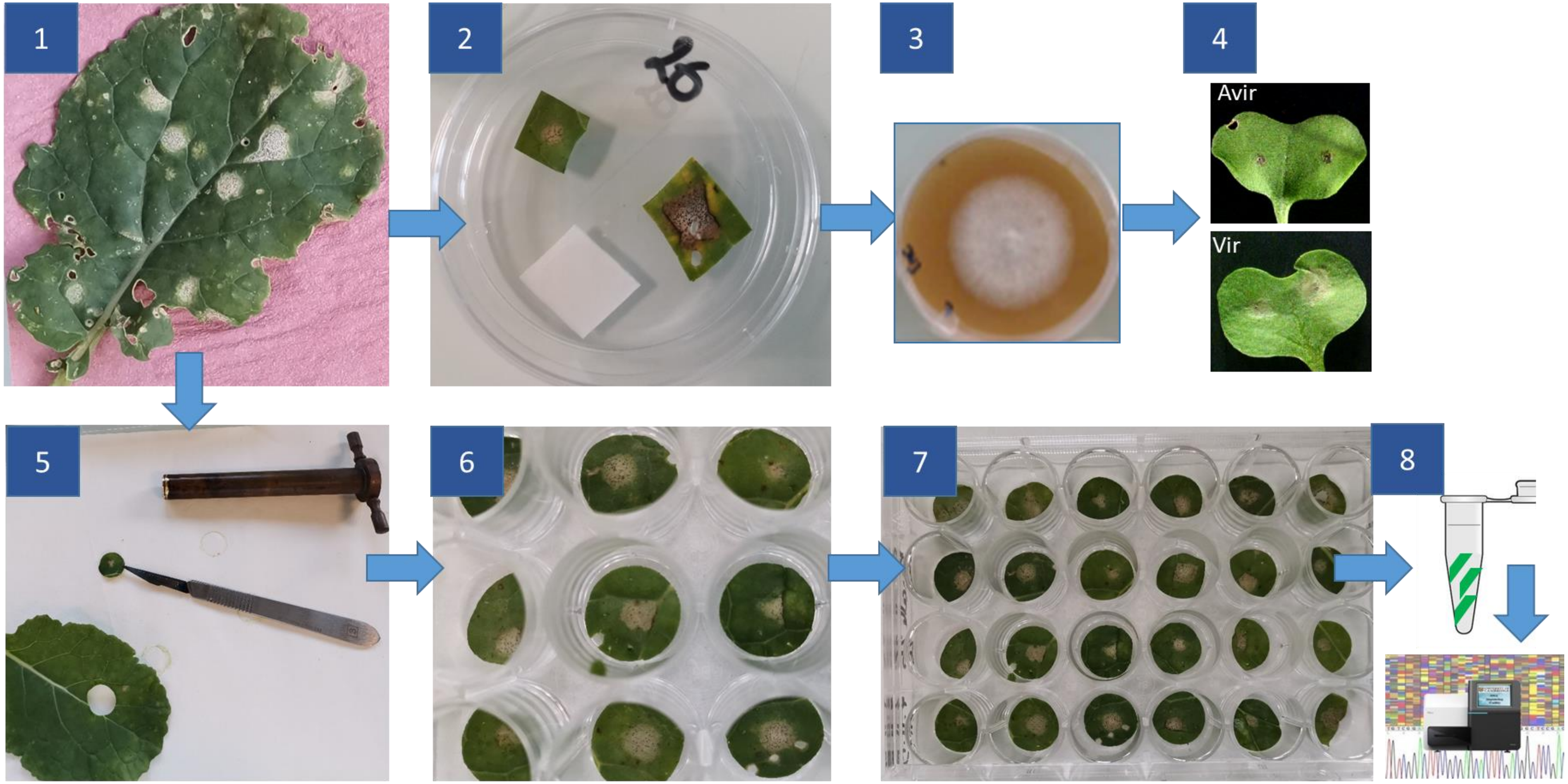

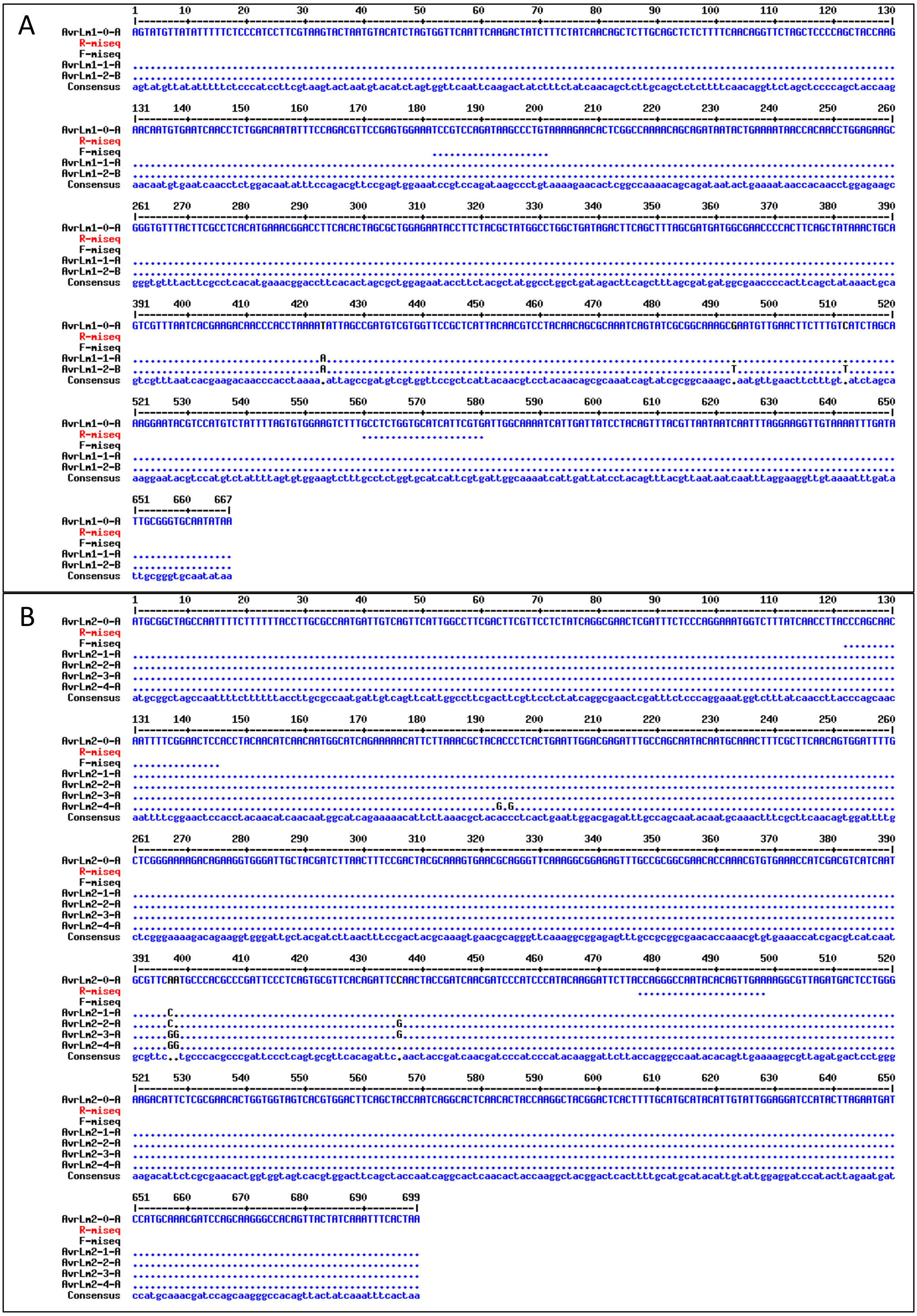
A
AvrLm1-0-A
R-miseq
F-miseq
AvrLm1-1-A
AvrLm1-2-B
Consensus
B
AvrLm2-0-A
R-miseq
F-miseq
AvrLm2-1-A
AvrLm2-2-A
AvrLm2-3-A
AvrLm2-4-A
Consensus

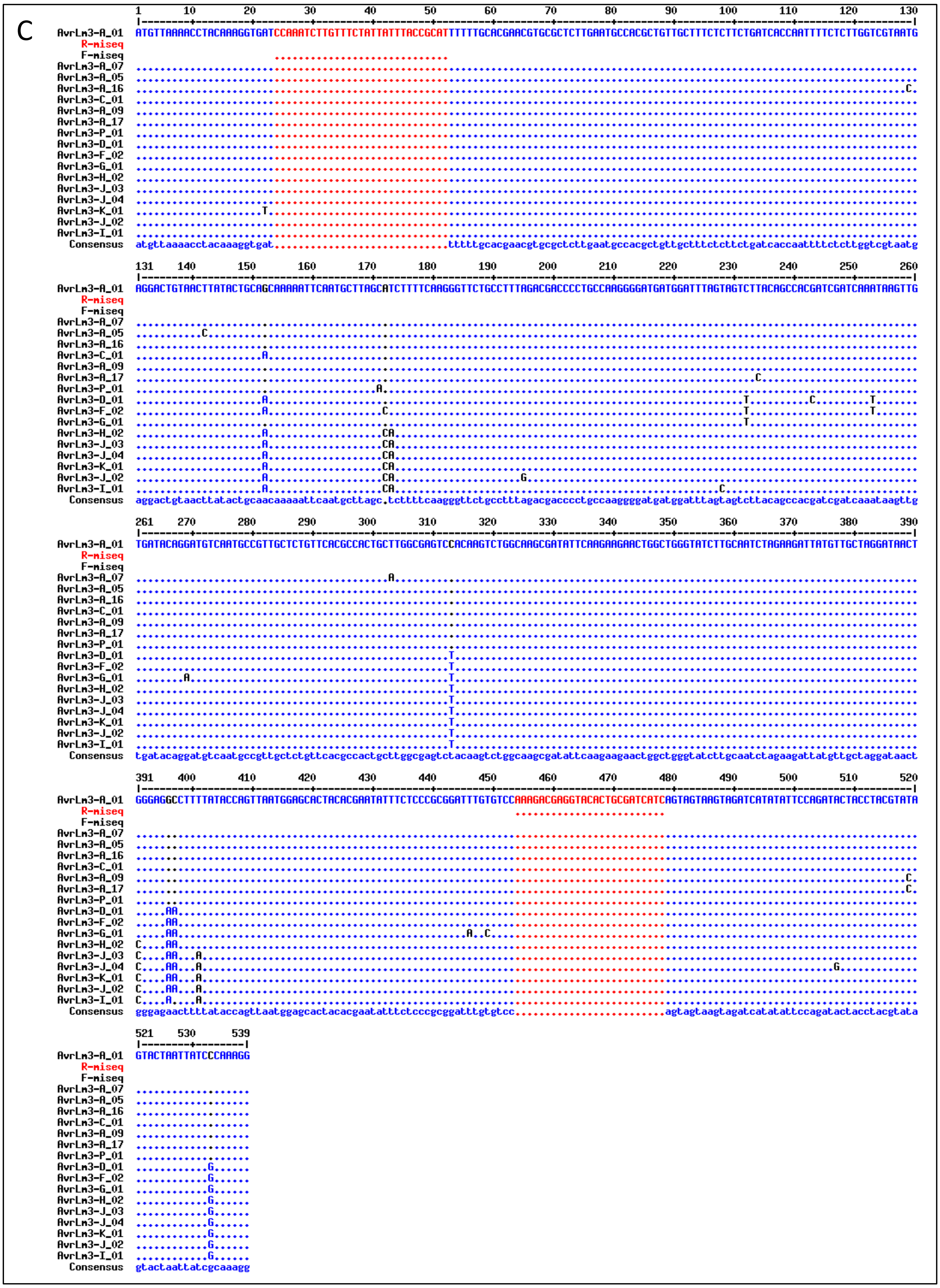
C
AvrLm3-A_01
R-miseq
F-miseq
AvrLm3-A_07
AvrLm3-A_05
AvrLm3-A_16
AvrLm3-C_01
AvrLm3-A_09
AvrLm3-A_17
AvrLm3-P_01
AvrLm3-D_01
AvrLm3-F_02
AvrLm3-G_01
AvrLm3-H_02
AvrLm3-J_03
AvrLm3-J_04
AvrLm3-K_01
AvrLm3-J_02
AvrLm3-I_01
Consensus
1 10 20 30 40 50 60 70 80 90 100 110 120 130
131 140 150 160 170 180 190 200 210 220 230 240 250 260
261 270 280 290 300 310 320 330 340 350 360 370 380 390

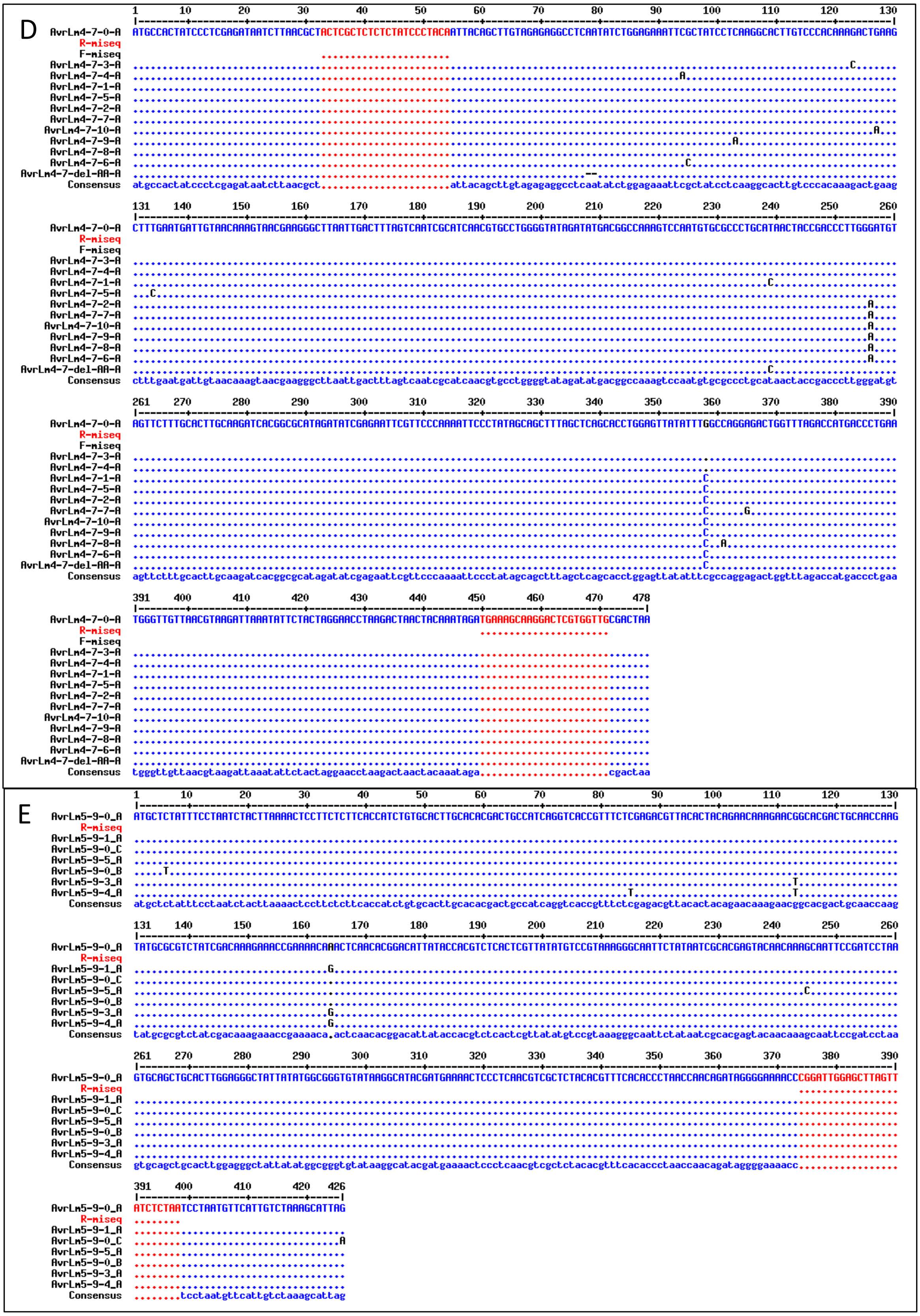
D
AvrLm4-7-0-A
R-miseq
F-miseq
AvrLm4-7-3-A
AvrLm4-7-4-A
AvrLm4-7-1-A
AvrLm4-7-5-A
AvrLm4-7-2-A
AvrLm4-7-7-A
AvrLm4-7-10-A
AvrLm4-7-9-A
AvrLm4-7-8-A
AvrLm4-7-6-A
AvrLm4-7-del-AA-A
Consensus
E
AvrLm5-9-0_A
R-miseq
AvrLm5-9-1_A
AvrLm5-9-0_C
AvrLm5-9-5_A
AvrLm5-9-0_B
AvrLm5-9-3_A
AvrLm5-9-4_A
Consensus

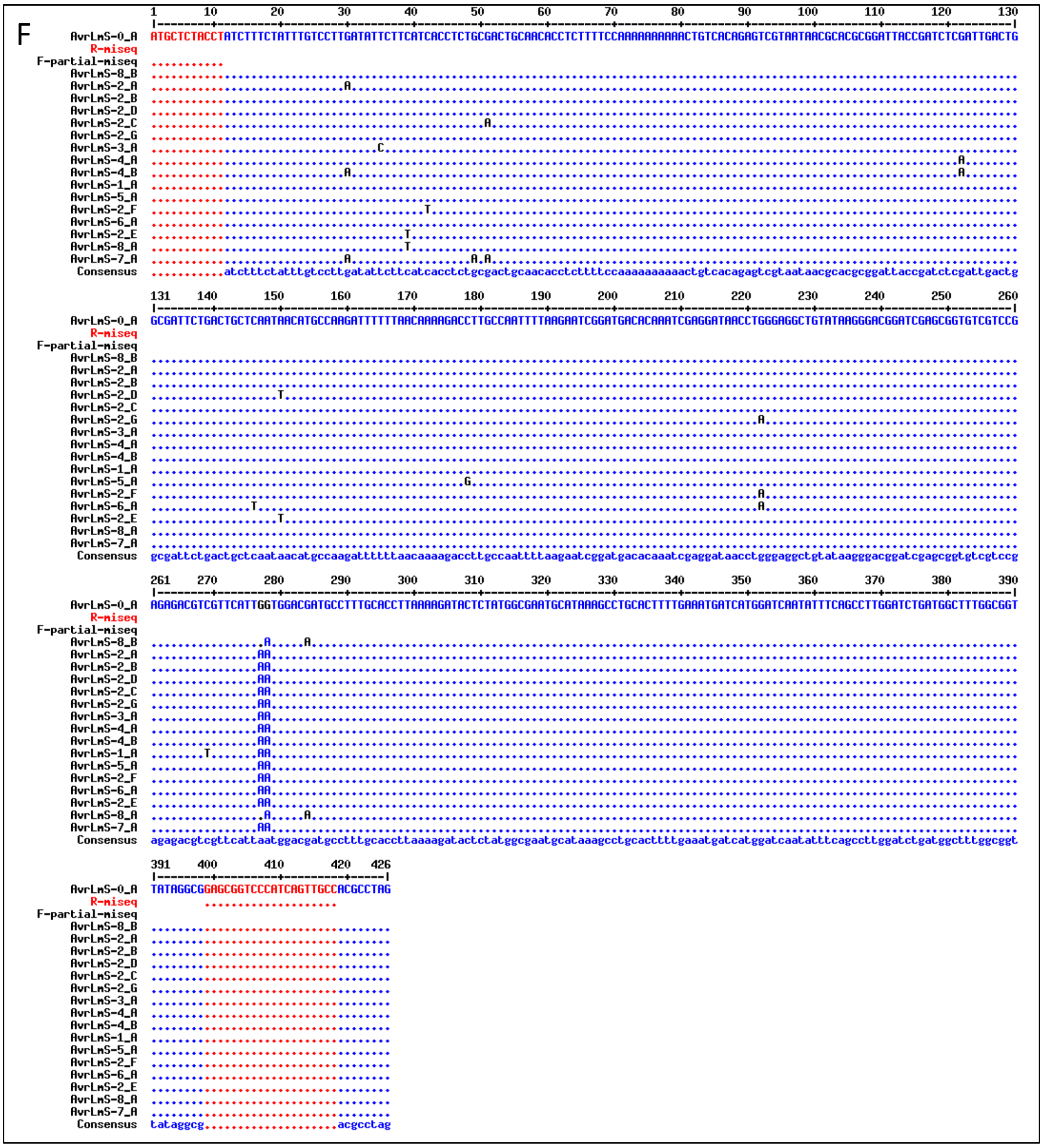
F
AvrLmS-0_A
R-miseq
F-partial-miseq
AvrLmS-8_B
AvrLmS-2_A
AvrLmS-2_B
AvrLmS-2_D
AvrLmS-2_C
AvrLmS-2_G
AvrLmS-3_A
AvrLmS-4_A
AvrLmS-4_B
AvrLmS-1_A
AvrLmS-5_A
AvrLmS-2_F
AvrLmS-6_A
AvrLmS-2_E
AvrLmS-8_A
AvrLmS-7_A
Consensus

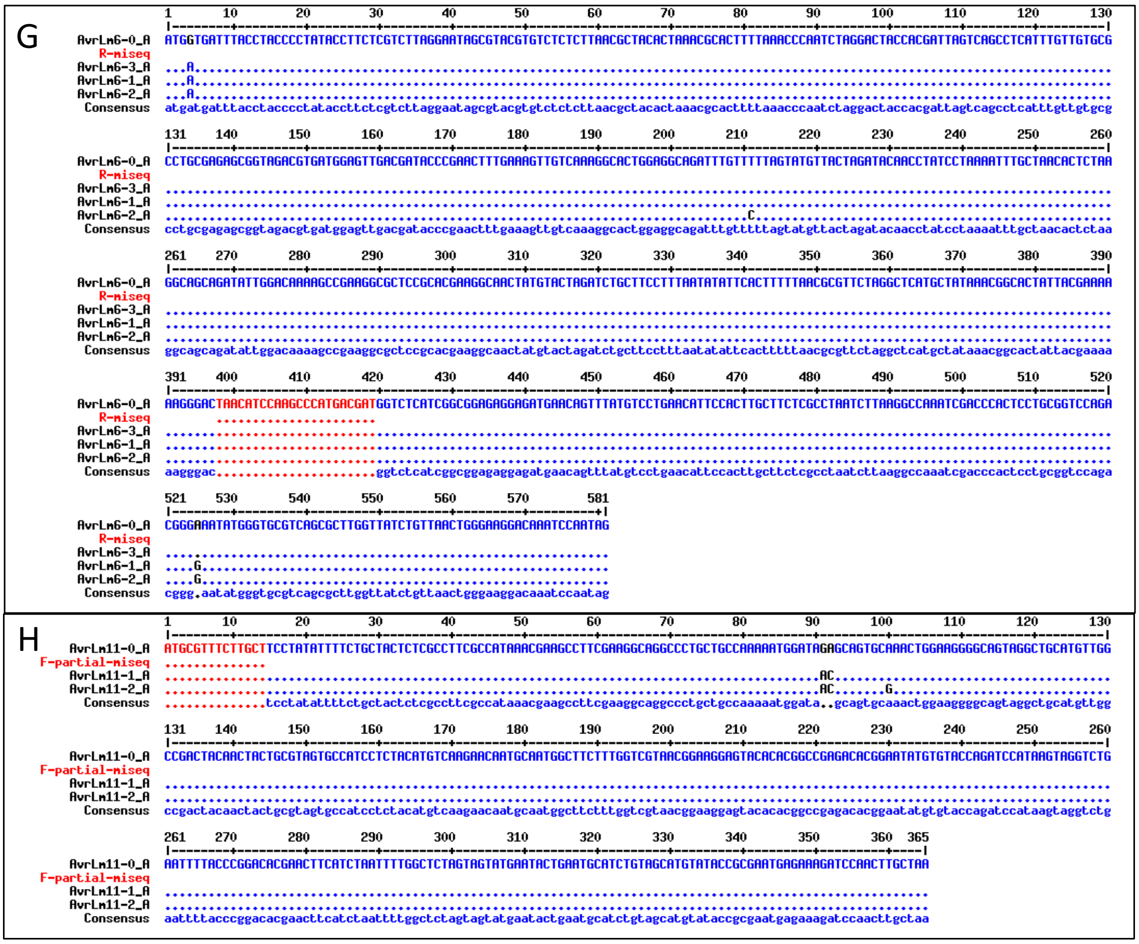

**Supplementary Figure S2. Alignment of known alleles of eight *L. maculans* avirulence genes with positions of forward and reverse MiSeq primers.** A, *AvrLm1*; B, *AvrLm2*; C, *AvrLm3*; D, *AvrLm4-7*; E, *AvrLm5-9*; F, *AvrLmS-Lep2*; G, *AvrLm6*; H, *AvrLm11*. The F-primers are located before the start for *AvrLm5-9* and *AvrLm6* and the R-primer is after the stop for *AvrLm11*.

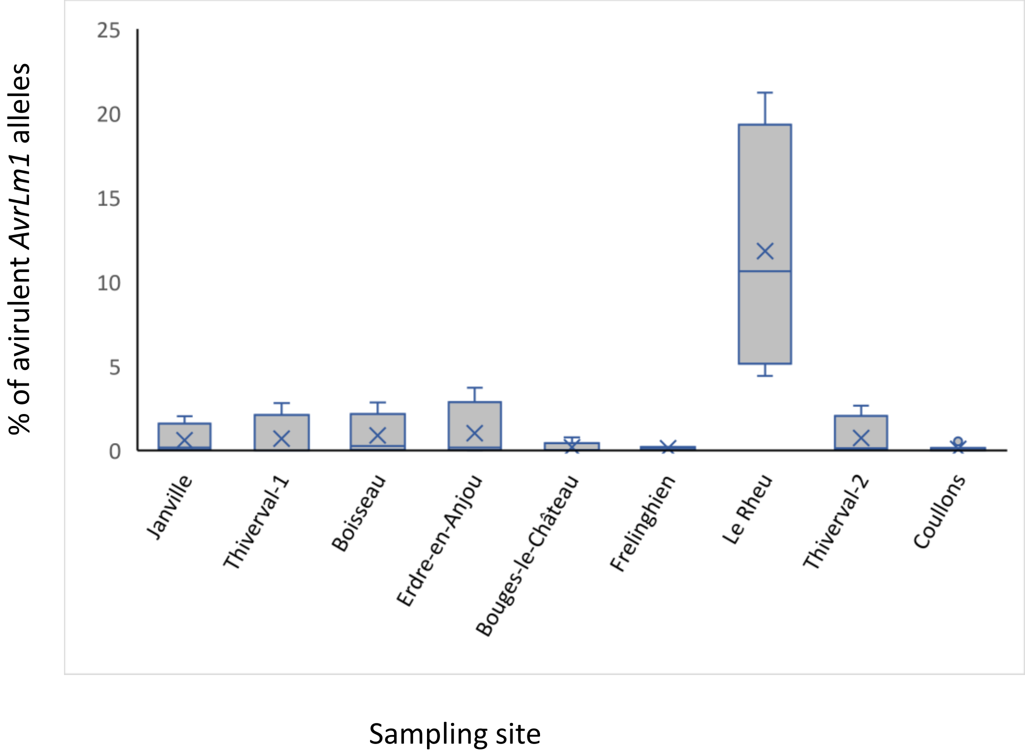


**Supplementary Figure S3**. Box plot of the frequency per sampling site of avirulent alleles for *AvrLm1,* as estimated by Miseq sequencing in 42 pools of leaf spots

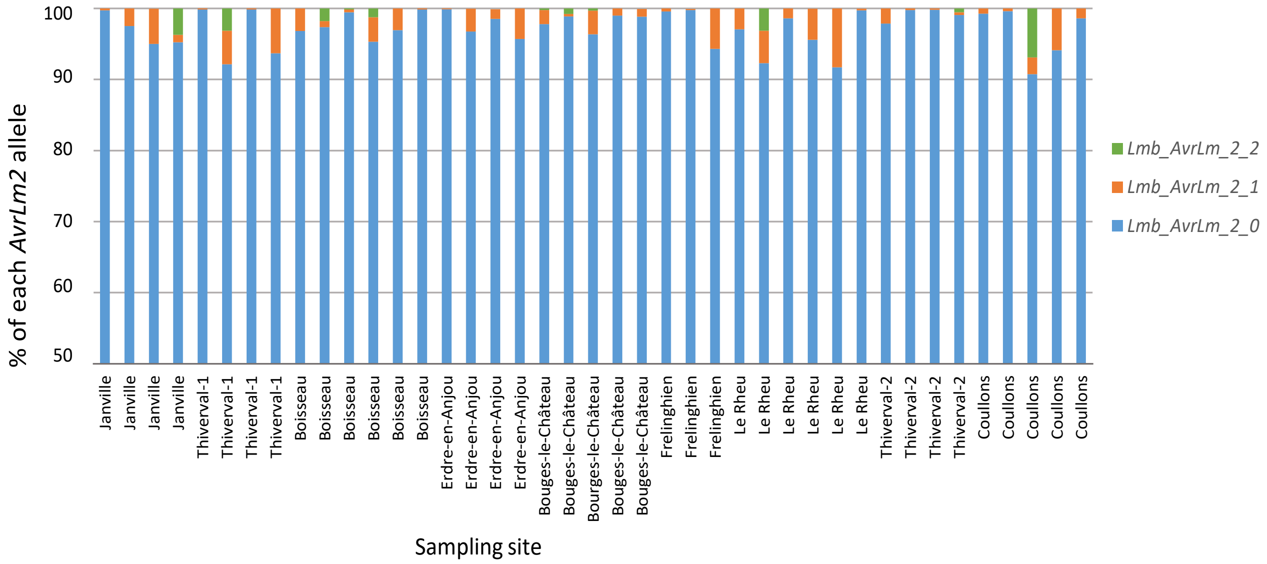


**Supplementary Figure S4. Proportions of the three *AvrLm2* alleles detected in 42 pools of leaf spots from nine fields**

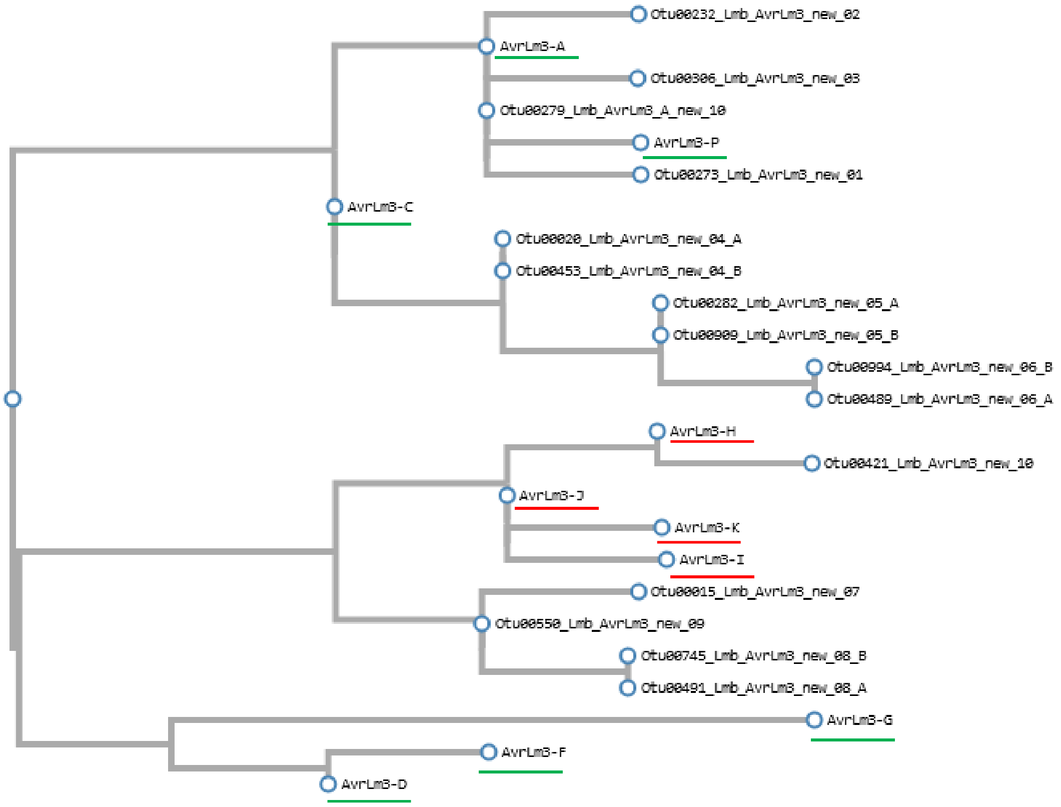


**Supplementary Figure S5 Maximum likelihood tree of all *AvrLm3* isoforms identified here.** Isoforms underlined in green are known avirulent isoforms while those underlined in red are known virulent isoforms.s.

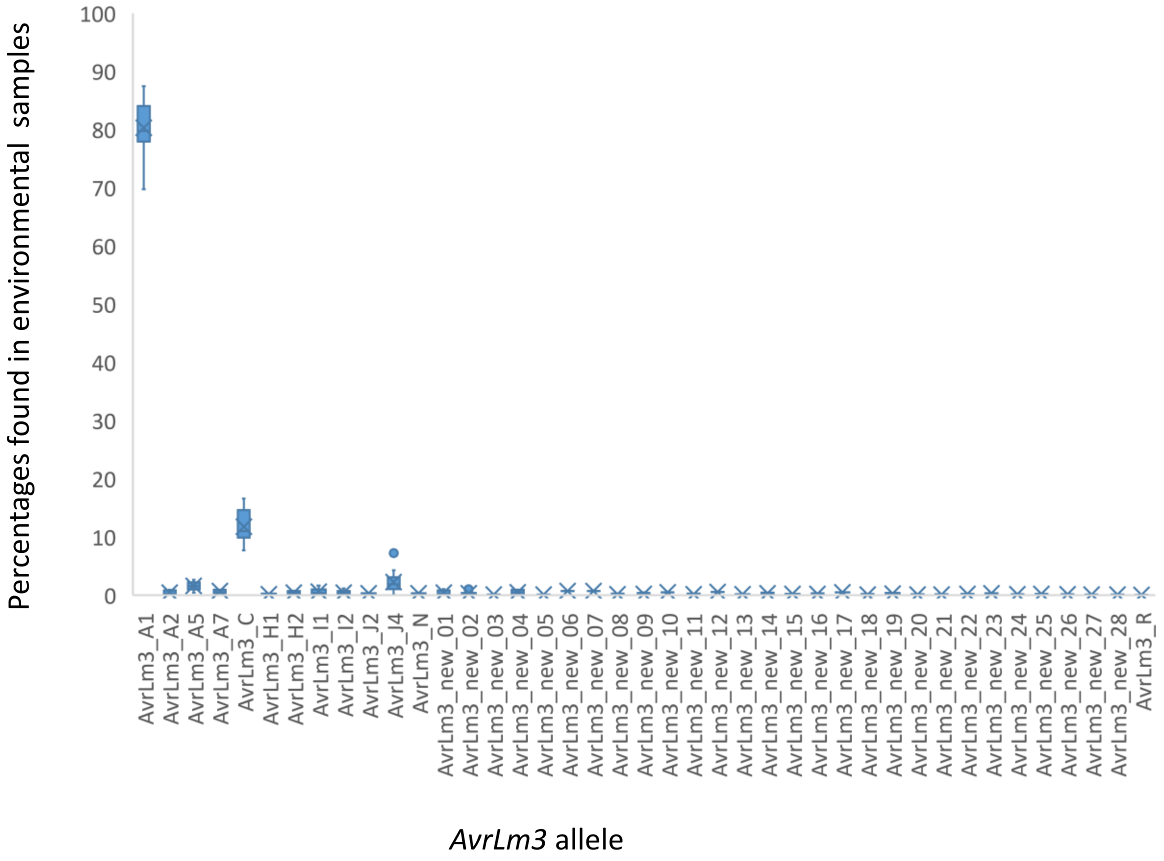


**Supplementary Figure S6. Percentages of the 41 *AvrLm3* alleles as determined following multiplex-PCR and MiSeq sequencing of 42 environmental samples**

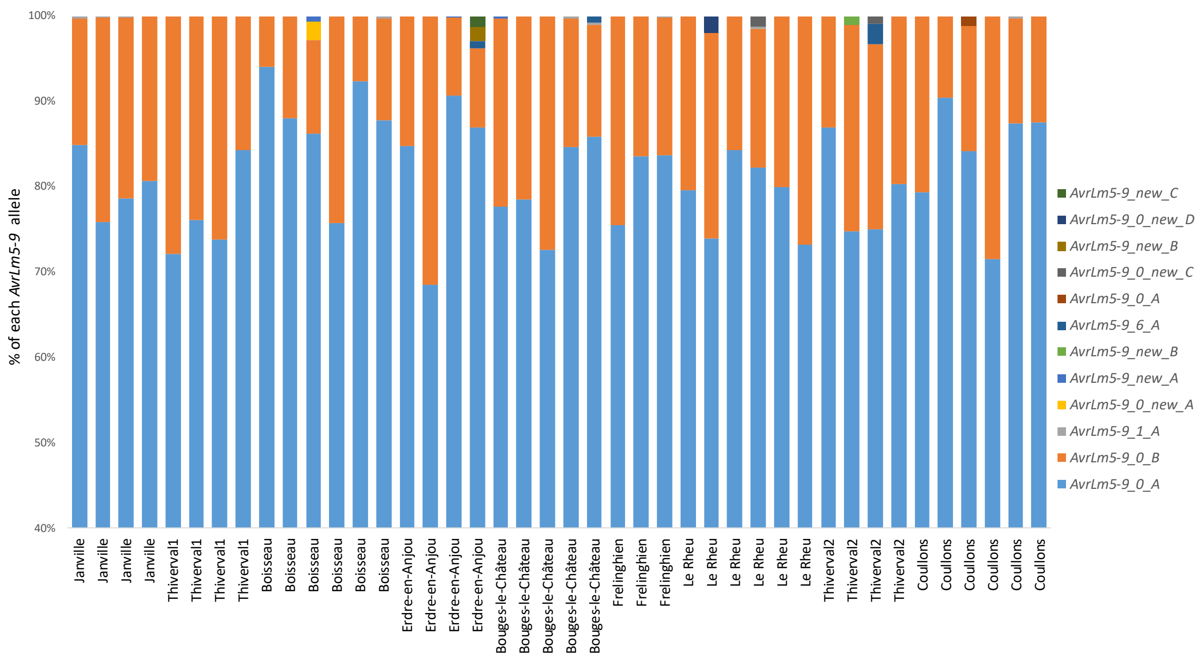


**Supplementary Figure S7. Proportions of *AvrLm5-9* alleles detected in 42 pools of leaf spots from nine fields**

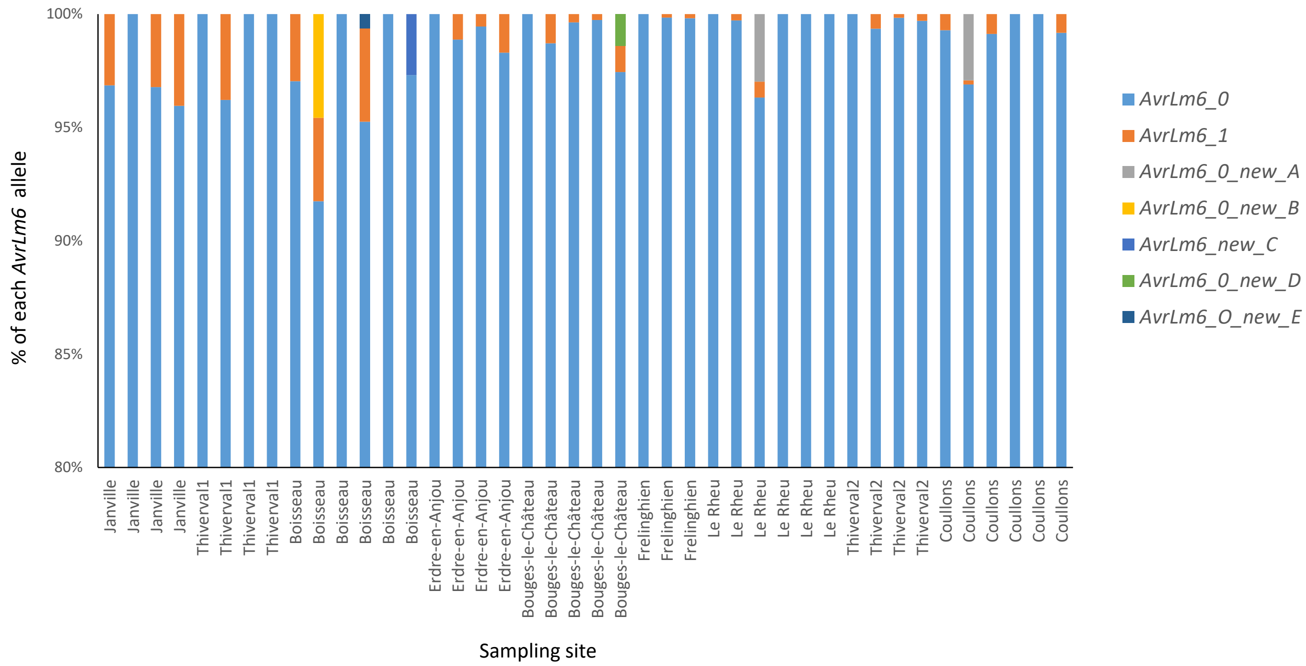


**Supplementary Figure S8. Proportions of *AvrLm6* alleles detected in 42 pools of leaf spots from nine fields**

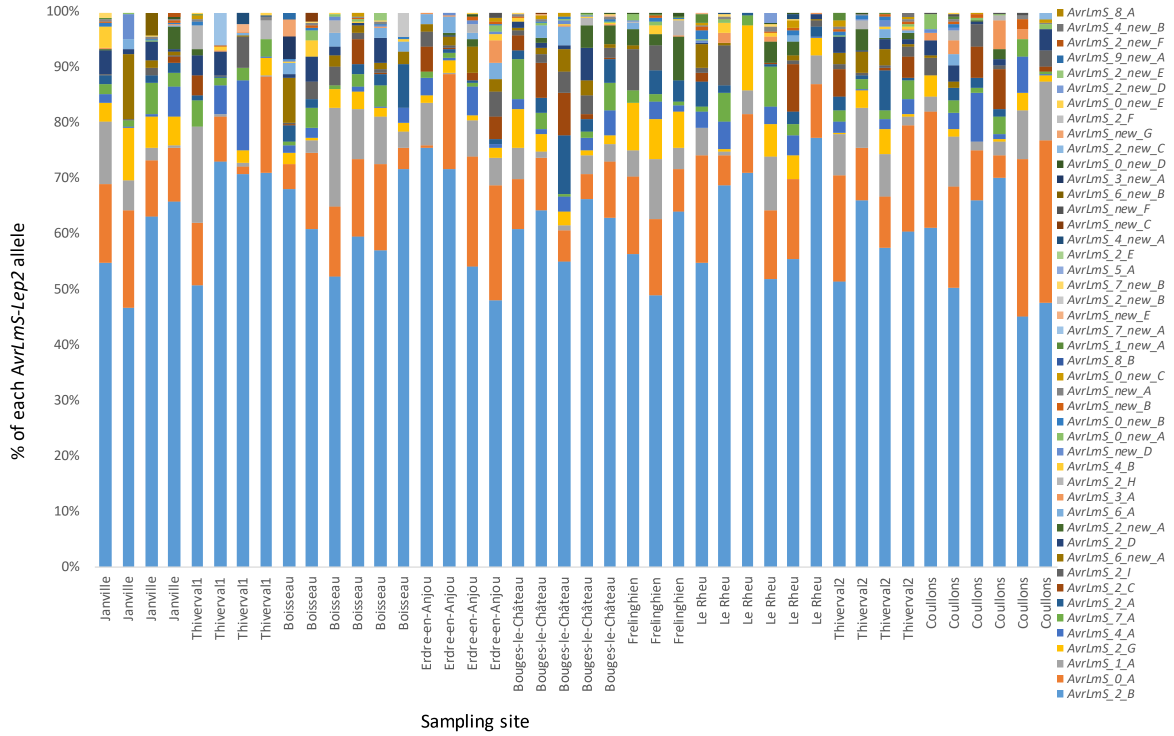


**Supplementary Figure S9. Proportions of *AvrLmS-Lep2* alleles detected in 42 pools of leaf spots from nine fields.**

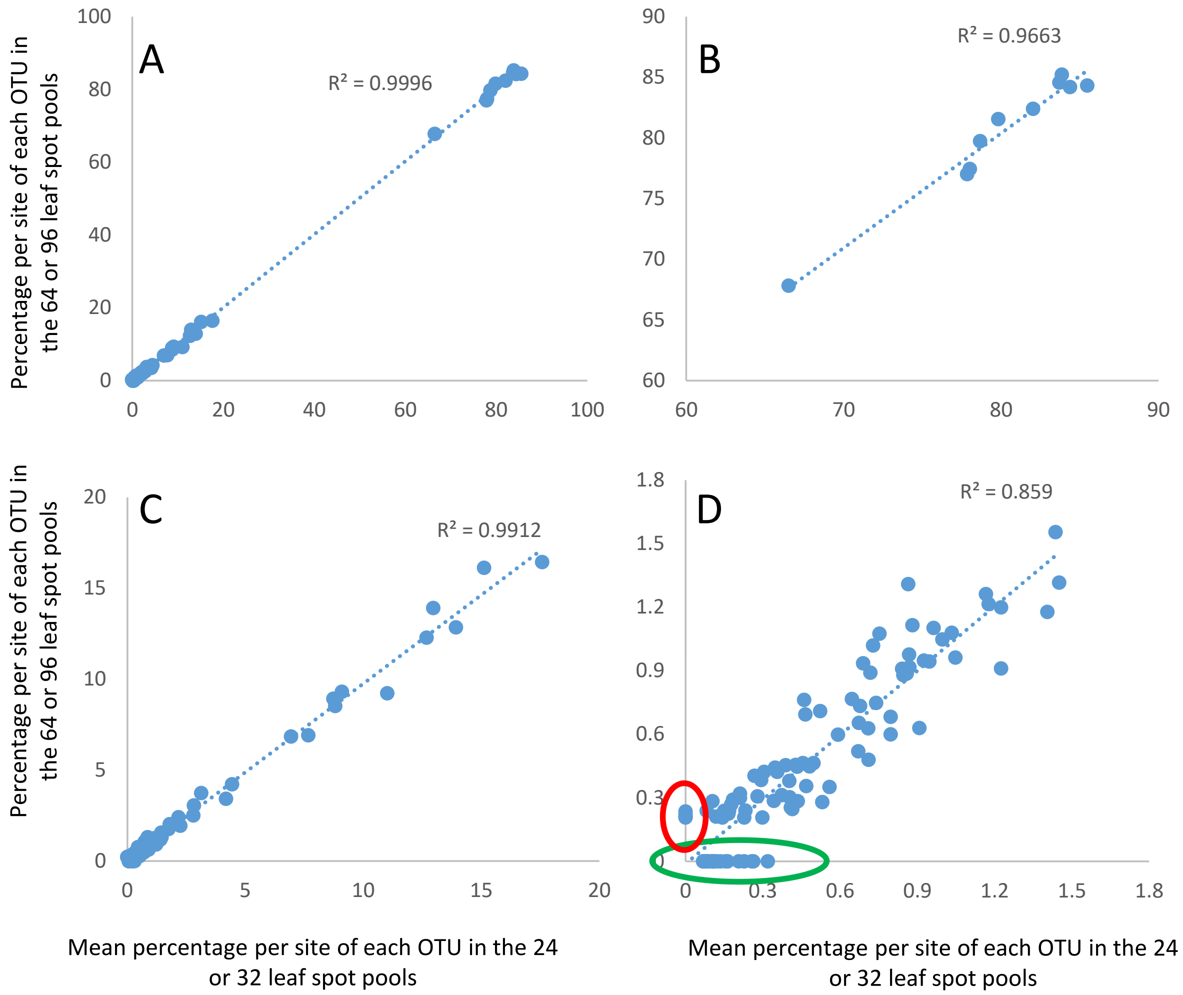


**Supplementary Figure S10. Relationships between percentages of each *AvrLm3* OTU per site estimated in 32 leaf spots or from the "pools of pools" (64 or 96 leaf spots). Each dot corresponds to the percentage of one OTU in one site.** A, all data; B, C and D, focus on high percentages (>60%), low percentages (<20%) or very low percentages (<2%). The green circle highlights OTUs that are detected at low frequencies in 24 or 32 leaf spot pools but not in 64 or 96 leaf spots. The red circle highlights OTUs that are not detected in pools of 32 leaf spots but found in pools of 96, thus probably corresponding to sequencing artefacts.

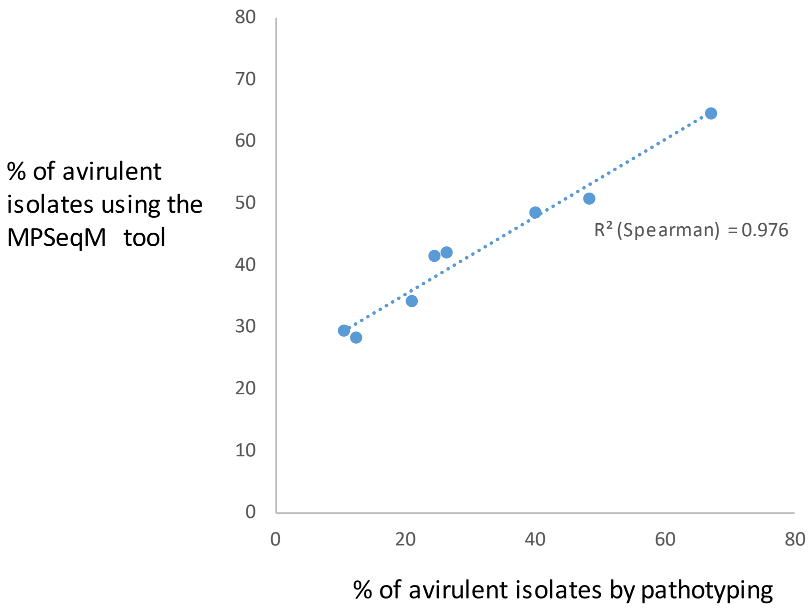


**Supplementary Figure S11. Correlation between estimates of percentage of avirulent isolates on *Rlm7* by pathotyping and with the MPSeqM tool.** Each dot corresponds to one geographical location. Samplings were done at the same place and same time but on different leaves. Pathotyping was described in Balesdent et al. (2023) and was done using around 80 isolates per site, while the genotyping was done on 3 to 6 pools of 32 leaf spots per site. The Spearman correlation coefficient is indicated.

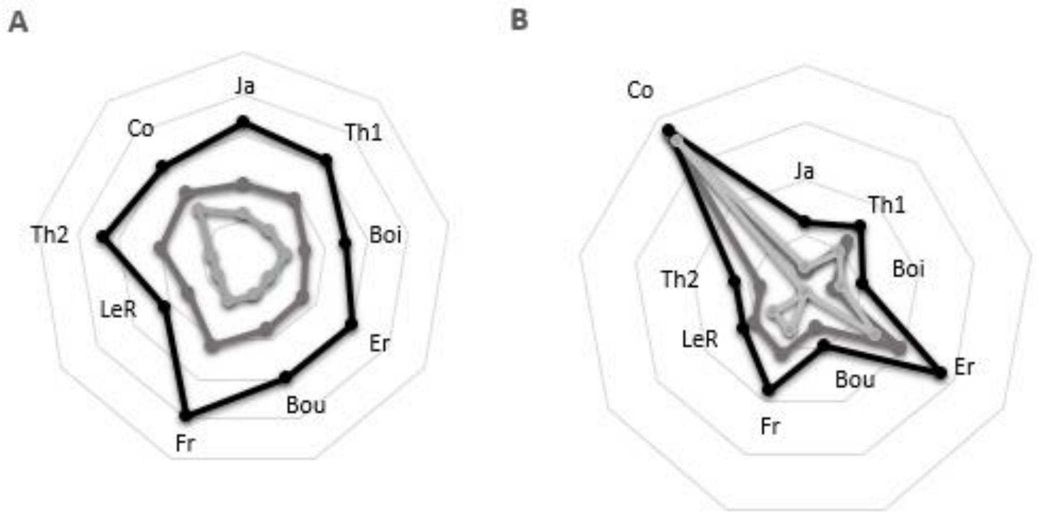


**Supplementary Figure S12. Relative *AvrLm* allele diversity per site as described with the MpSeqM tool.** A, diversity for all genes; B, diversity for *AvrLm11* only. The radar diagrams represent, for each site, the relative proportion of alleles (no. of alleles at this site over total no. of alleles detected in the survey) averaged by the relative proportion of leaf spots analysed at this site. Dark line, overall diversity; dark grey line, diversity for "new" alleles; pale grey line, diversity for site-specific new alleles. Sites as follows: Co, Coullons; Ja, Janville; Th1, Thiverval-Grignon field 1; Th2, Thiverval-Grignon field 2; Boi, Boisseau; Er, Erdre-en-Anjou; Bou, Bouges-le-Château; Fr, Frelinghein; LeR, Le Rheu

**Supplementary Text T1. Parameters used for the Mothur pipeline.**

**Pipeline according to Mothur MiSeq SOP**

### A. Parameter and process used

A.1 Tool Pear on Galaxy (https://vm-galaxy-prod.toulouse.inrae.fr/galaxy_main): Pairing reads R1 and R2 from fastq files

Minimum overlap size : 40
Maximum possible length of the assembled sequences : 460
Minimum possible length of the assembled sequences : 200
Minimum length of reads after trimming the low quality part : 100
Maximal proportion of uncalled bases in a read : 0

A.2 Search for primers to split dataset gene by gene : (100% primer identity)

A.3 Mothur analysis for each gene independently:

Analysis steps to be performed:

- Quality trim
- Group curated files
- Count unique sequences
- Search for chimera
- Classify against in house database
- Distance matrix calculation
- Clustering
- Create database

Specific parameters used for each gene at some steps are summarized in the table below:

| | | Actin | AvrLm1 | AvrLm2 | AvrLm3 | AvrLm4-7 | AvrLm5-9 | AvrLm6 | AvrLm11 | AvrLmS |
|---|---|---|---|---|---|---|---|---|---|---|
| B. 1) trim.seqs | maxambig | 0 | 0 | 0 | 0 | 0 | 0 | 0 | 0 | 0 |
| | maxhomop | 15 | 10 | 10 | 10 | 10 | 10 | 10 | 10 | 14 |
| | minlength | 150 | 350 | 325 | 390 | 385 | 375 | 395 | 390 | 380 |
| | qwindowaverage | 28 | 28 | 28 | 28 | 28 | 28 | 28 | 28 | **24** |
| | qwindowsize | 25 | 25 | 25 | 25 | 25 | 25 | 25 | 25 | 25 |
| B. 3) split.abund (nb min. reads by otu) | cutoff | 2 | 2 | 2 | 2 | 2 | 2 | 2 | 2 | 2 |
| B. 5) classify.seqs | iters | 1000 | 1000 | 1000 | 1000 | 1000 | 1000 | 1000 | 1000 | 1000 |
| B. 8) get.oturep(homolgy and longueur 100%) | | unique | unique | unique | unique | unique | unique | unique | unique | unique |

### B. Command-line : example for *AvrLm1* analysis

B.1 Quality and trim of the reads for each file (3 files as example A, B, and C)

*fastq.info(fastq=**A**.fastq)*
*fastq.info(fastq=**B**.fastq)*
*fastq.info(fastq=**C**.fastq)*
*trim.seqs(fasta=**A**.fasta, oligos=AvrLm1.oligos, qfile=**A**.qual, maxambig=0, maxhomop=10, minlength=350, qwindowaverage=28, qwindowsize=25)*
*trim.seqs(fasta=**B**.fasta, oligos=AvrLm1.oligos, qfile=**B**.qual, maxambig=0, maxhomop=10, minlength=350, qwindowaverage=28, qwindowsize=25)*

*trim.seqs(fasta=**C**.fasta, oligos=AvrLm1.oligos, qfile=**C**.qual, maxambig=0, maxhomop=10, minlength=350, qwindowaverage=28, qwindowsize=25)*

B.2 Group the curated reads of the different files in one file

*make.group(fasta=**A**.trim.fasta-**B**.trim.fasta-**C**.trim.fasta, groups=**A**-**B**-**C**, output=merge.groups)*
*merge.files(input=**A**.trim.fasta-**B**.trim.fasta-**C**.trim.fasta, output=merge.AVRLM1.fasta)*

B.3 Search and count unique sequences

*unique.seqs(fasta=merge.AVRLM1.fasta)*
*count.seqs(name=merge.AVRLM1.names, group=merge.groups)*
*summary.seqs(fasta=**A**.trim.fasta)*
*summary.seqs(fasta=**B**.trim.fasta)*
*summary.seqs(fasta=**C**.trim.fasta)*
*summary.seqs(fasta=merge.AVRLM1.fasta)*
*summary.seqs(fasta=merge.AVRLM1.unique.fasta)*
*split.abund(fasta=merge.AVRLM1.unique.fasta, name=merge.AVRLM1.names, cutoff=2, accnos=true)*

B.4 Search for chimera

*chimera.uchime(fasta=merge.AVRLM1.unique.abund.fasta, name=merge.AVRLM1.abund.names, reference=self)*
*remove.seqs(accnos=merge.AVRLM1.unique.abund.denovo.uchime.accnos, fasta=merge.AVRLM1.unique.abund.fasta, name=merge.AVRLM1.abund.names)*

B.5 Classify against database AVRLM reference file (**Lmb_avrlm_act.reference.txt; Lmb_avrlm_act.taxonomy.txt**)

*classify.seqs(fasta=merge.AVRLM1.unique.abund.pick.fasta, template=Lmb_avrlm_act.reference.txt, taxonomy=Lmb_avrlm_act.taxonomy.txt, iters=1000, name=merge.groups)*

B.6 Distance matrix calculation

*pairwise.seqs(fasta=merge.AVRLM1.unique.abund.pick.fasta, processors=60)*

B.7 Clustering

*cluster(column=merge.AVRLM1.unique.abund.pick.dist, name=merge.AVRLM1.abund.pick.names)*

B.8 Create database

*get.oturep(column=merge.AVRLM1unique.abund.pick.dist, name=merge.AVRLM1.abund.pick.names, fasta=merge.AVRLM1.unique.abund.pick.fasta, list=merge.AVRLM1.unique.abund.pick.an.list, label=unique)*
*classify.otu(taxonomy=merge.AVRLM1.unique.abund.pick.taxonomy.wang.taxonomy, name=merge.AVRLM1.abund.pick.names, list=merge.AVRLM1.unique.abund.pick.an.list, reftaxonomy=Lmb_avrlm_act.taxonomy.txt)*
*create.database(list=merge.AVRLM1.unique.abund.pick.an.list, label=unique, repfasta=merge.AVRLM1.unique.abund.pick.an.unique.rep.fasta, repname=merge.AVRLM1.unique.abund.pick.an.unique.rep.names, constaxonomy=merge.AVRLM1.unique.abund.pick.an.unique.cons.taxonomy, group=merge.groups)*
